# Compositional data modeling of high-dimensional single cell RNA-seq (CoDA-hd): its advantages over commonly used normalization approaches

**DOI:** 10.1101/2025.03.24.644852

**Authors:** Jinghan Huang, Phillip Sheung Chi Yam, KS Leung, Minghua Deng, Nelson LS Tang

**Affiliations:** Department of Chemical Pathology, Li Ka Shing Institute of Health Science, Faculty of Medicine, The Chinese University of Hong Kong, Hong Kong SAR, China; Department of Statistics, The Chinese University of Hong Kong, Hong Kong SAR, China; Department of Computer Science and Engineering (CSE), The Chinese University of Hong Kong, Hong Kong SAR, China; Cytomics Limited, Hong Kong Science Park, Hong Kong SAR, China; School of Mathematical Sciences, Peking University, Beijing, China; Hong Kong Branch of CAS Center for Excellence in Animal Evolution and Genetics and KIZ/CUHK Joint Laboratory of Bioresources and Molecular Research in Common Diseases, Hong Kong SAR, China; Functional Genomics and Biostatistical Computing Laboratory, CUHK Shenzhen Research Institute, Shenzhen, China

**Author notes:** **Correspondence:** Nelson L.S. TANG, MBChB, MD, FRCPA, And Jinghan Huang, Department of Chemical Pathology and Li Ka Shing Institute of Health Sciences (LiHS), Faculty of Medicine, The Chinese University of Hong Kong.

**Keywords:** Single cell RNA-seq, Compositional data analysis, Clustering, Trajectory inference

## Abstract

Compositional data analysis (CoDA) is an emerging statistical framework and has been extended to microbiome, bulk RNA-seq, and cell type proportions in single-cell RNA-seq (scRNA-seq) (with 50-200 components). Here, we explore the high-dimensional application of CoDA (CoDA-hd) and its various log-ratio (LR) transformations to raw count matrices of scRNA-seq which has over 20,000 components (e.g., protein coding genes). scRNA-seq matrices are typically sparse and high-dimensional. Common approaches of normalization such as log-normalization may lead to suspicious findings as previously shown for trajectory inference. Although RNA-seq is compositional data by nature, the geometry of CoDA in high-dimensional simplex is not compatible with most downstream analyses of scRNA-seq which are based on Euclidean space. In this study, we explored if CoDA is adaptable with scRNA-seq in various downstream applications. Specifically, we attempted to study: (1) CoDA adaptability to scRNA-seq; (2) handling of zero data: prior-log-normalized, imputation or with specific count addition; (3) transformation to Euclidean space and compatibility with downstream analyses. Our results suggested that (1) the innovative count addition schemes (e.g., SGM) enable the application of CoDA to high dimensional sparse data (i.e., scRNA-seq); (2) log-normalized data could be transformed to CoDA LR transformation as an approximation; (3) CoDA LR transformations such as count-added centered-log-ratio (CLR) had some advantages in dimension reduction visualization, clustering, and trajectory inference in the tested real & simulated datasets. CLR provided more decent and separated clusters in dimension reductions, improved the Slingshot trajectory inference, and eliminated the suspicious trajectory that is probably caused by the dropouts. We therefore concluded that CoDA may be a preferred scale-free model to handle scRNA-seq data for these downstream applications. Additionally, an R package ‘CoDAhd’ was developed for conducting CoDA LR transformations for high dimensional scRNA-seq data. The code for implementing CoDA-hd and some example datasets were placed at https://github.com/GO3295/CoDAhd.

## Introduction

Single cell RNA sequencing (scRNA-seq) has been widely used in biological and biomedical studies for exploring transcriptomic profiling and differences in proportions of various cell populations. In scRNA-seq data analysis, the first step is the normalization of raw counts of transcript abundances (TA) of genes so that total counts of TA are comparable across individual cells. The conventional normalization method is the log-normalization as described and reviewed previously^1^. Subsequently, researchers developed SCTransform which applied regularized negative binomial regression^2^ to pre-process the data^3^. Typically, scRNA-seq data matrix is ∼20,000 genes × thousands to millions cells (20,000× 100,000+) that are captured for analysis and the gene expressions of every cell are represented as the counts in the matrix.

The dropout problem is an important but also a common artifact in scRNA-seq. Due to poorly understood experimental and technical factors, TA counts of some random collections of genes are zero. This phenomenon is not limited to low-expression genes, as housekeeping genes may also have zero TA counts in some cells. Although scRNA-seq data is CoDA by nature, there was no previous attempts of applying CoDA due to these difficulties (i.e., extreme high dimension and extremely sparse matrix). Most conventional normalization methods are not robust to such randomly assigned missing data. For example, PCA/UMAP using conventional methods may lead to suspicious findings due to the ‘dropouts’ problem (e.g., a controversial differentiation trajectory from plasmablasts to developing neutrophils using existing normalization methods, which was not plausible according to our current understandings^4–6^). Therefore, we are exploring the utility of other statistical models and methods of data representation which are more robust for these properties and limitations of scRNA-seq data.

Here, we attempted for the first time the application of compositional data analysis (CoDA) in the analysis of scRNA-seq. As an emerging school of statistical data analysis, CoDA framework was first proposed by John Aitchison in the 80’s when it was applied to geochemical data and more recently to microbiome data (**Figure S1**). The core difference between CoDA and the conventional approach of scRNA-seq analysis is that, in order to transform data projection from the compositional simplex geometry to Euclidean space, CoDA handles data deliberately as the log-ratios (LR) between the components. In contrast, the existing scRNAseq analysis treats the TA data as real numbers in log space. Treating TA data by LR instead of real number should gain benefits with three key CoDA properties, which are scale invariance, sub-compositional coherence, and permutation invariance^7^. Scale invariance states that any scale (e.g., multiplied by a scale factor) of the original data will have no effect since compositional data only carries relative information. Sub-compositional coherence means that results obtained from a sub-composition (i.e., subset of data) will remain the same as in the composition. Permutation invariance is straightforward and states that the results do not depend on the order that the parts appear in a composition^8^. Another benefit of CoDA log-ratios is that it reduces the data skewness and make the data more balance for downstream analyses. These properties of CoDA make it an appropriate model for scRNA-seq data.

Theoretically, CoDA has great potential to be used in high-throughput sequencing (HTS) data. Its adaptability and application to microbiome data and bulk RNA sequencing data have been demonstrated. For microbiome data, bacterium proportions (**Figure 1A, Bacterium**) in a stool sample are compositional in nature and CoDA is readily applied to them. Recently, it is proposed that CoDA could be used to analyze bulk RNA-seq data due to the compositional nature of the data^8–10^. Bulk RNA-seq summates TA of all cells present in one sample and has been frequently used to explore the TA difference between different conditions. Thus, the data matrix is much smaller than scRNA-seq and the statistical questions mainly focus on the difference between conditions. In fact, HTS (e.g., bulk or single cell RNA-seq) naturally generate relative information of feature abundance and thus has an upper limit of total number of reads. This property was due to the sequencer as it can only sequence a fixed number of nucleotide fragments. Therefore, a theoretical competitive situation may occur where the number of one transcript increases will actually decrease the number of all other transcripts observed^8^. Some HTS examples includes studies applying CoDA to analyze the meta-genomics datasets for various microbiome and demonstrating its great performance^11–15^. Besides, researchers have developed several R CoDA packages to perform the gene differential abundance analysis, conduct simulation study or study cell type proportions (**Figure 1A, scRNA-seq cell types**) ^15–18^, usually for data matrix of size up to 1,000 × 1,000. Various benchmarks for comparing different methods with CoDA were also conducted^8,12,19–22^. In CITE-seq, the size of the matrix of cell surface protein expression is usually ∼100 proteins × thousands to millions cells, and the data is often handled by CoDA centered-log-ratio (CLR) transformation^23^. Compared to these existing CoDA applications for biological data, treating genes as components in CoDA for scRNA-seq data faces some major challenges, e.g., it consists of thousands to millions of individual cells to handle and can result in 1000-fold increase in columns. As the sizes of the data matrices range from >10,000×thousands to millions cells, we applied CoDA and named it CoDA-high dimensional (CoDA-hd) and explore the adaptability and output of such applications (**Figure 1A, scRNA-seq Genes**).

**Figure 1.**
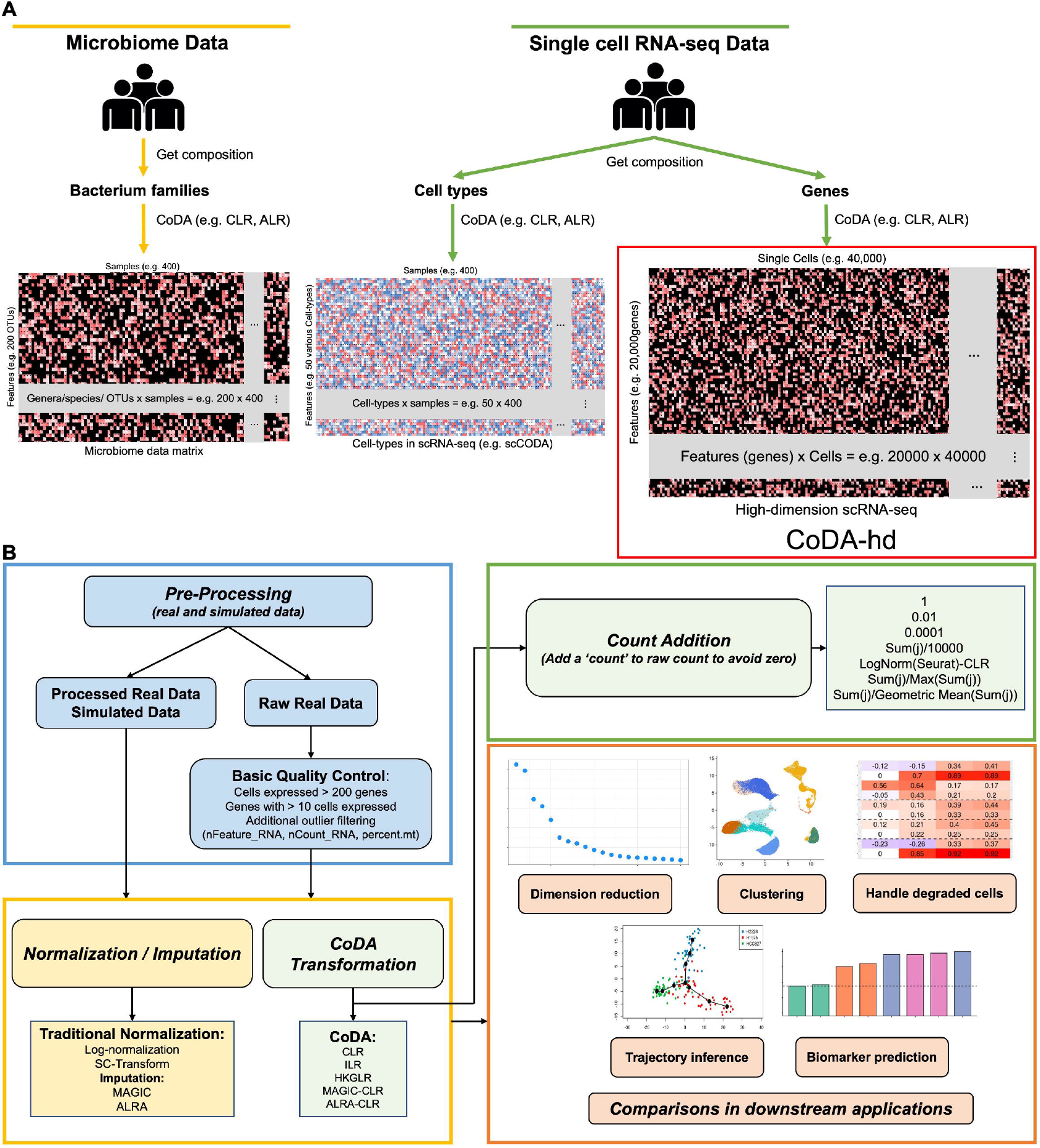
Concepts of compositional data analysis (CoDA) for different high-throughput sequencing data and the workflow in this study. (**A**) Current applications of compositional data analysis (CoDA) in different high-throughput sequencing data are illustrated. At present, CoDA only confined to hundreds of components or features. For example, there are hundreds of OTUs in Microbiome data and up to fifty cell type in cell count proportion data. Our study extends its use in much higher dimensional scRNA-seq data (up to tens of thousands of components) where each gene is treated as one component. (**B**) The workflow of analysis of twenty-nine real datasets (including a collection of 15 ‘gold standard’ trajectory datasets) and four simulated datasets (i.e., Splatter and SplatPop) which are pre-processed with quality control, normalization / imputation, or CoDA transformations. Performances of various CoDA transformations are evaluated by commonly used algorithms for downstream applications.

The scRNA-seq is very sparse: it has a large proportion of zeros and is possible to lead to false discovery or ambiguous conclusion as mentioned above^4^. How to handle these zeros, which is the key challenge in CoDA data transformations, was explored and compared in this study as well (**Figure 1B** and **Figure 2**). It has been previously mentioned that low counts matrices faced great challenges when processing with CoDA^22^. Indeed, previous research has demonstrated that applying a transformation for conceptual reasons does not necessarily translate into better downstream analysis results^3^.

**Figure 2.**
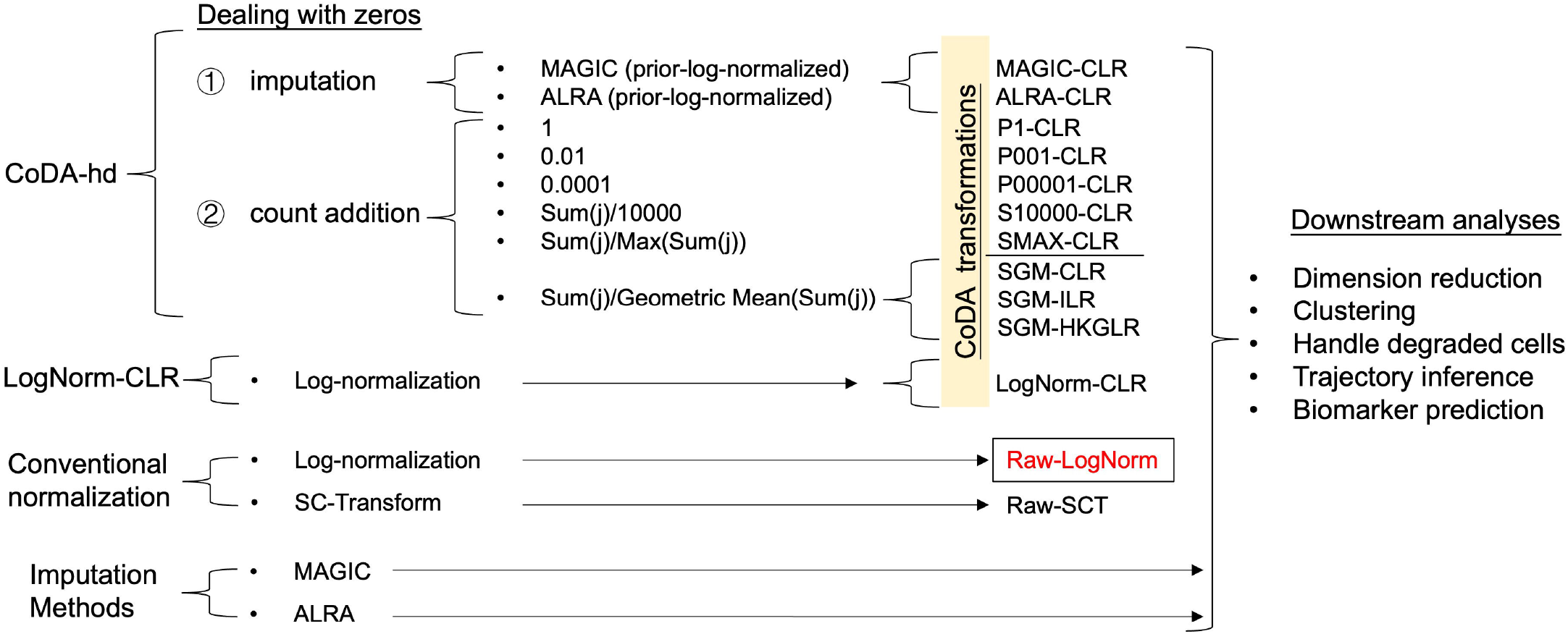
Details of the strategies of comparison among CoDA, conventional normalization and imputation methods, and overview of workflows in this study. Details of the framework in this study are presented. In CoDA-hd framework, two categories of methods (i.e., imputation and count addition) are applied to deal with zeros in scRNA-seq datasets before LR transformation by various log-ratio representations. In addition, conventional normalized data and imputed data are also forwarded for LR transformation. The effects with or without LR transformation were evaluated by five categories of downstream analyses (including dimension reduction, cell clustering, interference due to degraded cell, trajectory analysis and biomarker identification). Raw-LogNorm representing the typical Log-Normal transformation routine applied to scRNA-seq data by Seurat package is used as the baseline in all subsequent comparisons for CoDA performance.

Since the application of CoDA in scRNA-seq has not been explored, here we assess if CoDA as a normalization approach, which treats the TA counts of every single cell as compositional (**Figure 1A**), can be more robust and perform better in various scRNA-seq downstream applications (**Figure 1B, Figure 2** and **Table S1**). Motivated by the potential limitations of traditional log-normalization, we apply CoDA for both real and simulated scRNA-seq datasets and compare its performance with other methods.

## Methods

### Conventional normalization methods for scRNA-seq data

Existing normalization methods ignored the compositional nature of scRNA-seq data and just treat the dataset in Euclidean space. Furthermore, many data repositories only provide data after normalization. It has not been explored if such prior-normalized data matrices are compatible with subsequent CoDA transformation for LR analysis. The two common normalization methods (**Table S2, Group A**) are (1) log-normalization, by the function NormalizeData() and (2) SC-Transformation (regularized negative binomial regression) using SCTransform(). Both log-normalization and SC-Transform were implemented in Seurat^2^. Theoretically, data after log-normalization (default normalization in Seurat) could be transformed to CoDA LR transformation with scaling (as shown below).

### Transform scRNA-seq data to CoDA log-ratio (LR) transformations

In CoDA, it is assumed the real counts are not observed, and the only valid data is the composition or relative proportions between parts. Part is the terminology used in CoDA for components or variables, and genes in the context of scRNA-seq. The principal difference between CoDA and existing analysis methods is that data are represented ratios or log-ratios of parts in CoDA, while commonly used non-CoDA methods treated the data as real numbers. The key hurdle of using CoDA to analyze scRNA-seq data is its sparse data matrix. LR is required to transform the original CoDA data present in a simplex geometry compatible with most downstream analyses designated in Euclidean geometry^20^. Therefore, zero counts are incompatible with CoDA. Here, we explored two categories of methods to replace zero, (1) count addition and (2) imputation of missing data. A new method of count addition proposed here may be the most optimal method applicable to scRNA-seq. Furthermore, we compared our novel count addition CoDA approach with conventional normalization methods (log-normalization, SC-Transform), and imputation methods (MAGIC, ALRA) for scRNA-seq datasets.

#### 1. Handling of sparse matrix by count addition and CoDA LR transformations

In CoDA, each component (gene) represents a certain part taking up a particular percentage component of the whole specimen and all measured parts add up to a unit sum (e.g., 100%). For TA count data, TA of all genes summed up to 1 for every individual cell and each gene is represented as the proportional transcript abundance among transcripts of all genes expressed in that cell. This proportional abundance is further transformed by various LR methods for CoDA. To avoid zeros in LR process, we applied various commonly used count addition and a new proposal of count addition that could be more applicable to single cell HD matrix to the raw counts and evaluated their effects (**Figure 1B** and **Figure 2**).

Theoretically, CoDA LR can be applied after count-addition to the sparse and zero-inflated scRNA-seq data. This is also the widely used method in typical CoDA of standard data matrix of lower dimension.

a. Count addition by a constant value (constant value addition). Previous studies reported several ways to handle zeros for CoDA such as the addition of half minimum value to only those zeros (i.e., a fixed assumed value for counts below the detect limit)^7^. However, these methods did not work well for scRNA-seq data as shown in results. Specifically, a fixed constant of <u>1</u>, <u>0.01</u>, or <u>0.0001</u> was added to all raw counts in the whole matrix before closure. And the LR transformations (e.g., CLR) were performed on the proportions after closure:

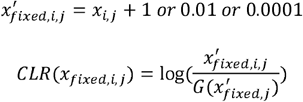 Where *x*_*i*,*j*_ is the raw count of the *i*gene and *j* cell. 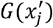 is the geometric mean of all genes of cell *j*. We discovered that the constant value addition method resulted in poor performance (see results). Therefore, other count addition methods were explored.
b. Count addition by a fraction with constant scaling denominator. Scaling to the same total count per cell (each column) in the experiment is a common practice in normalization by scaling, e.g., log-normalization (LogNorm). Here, the pseudocount (terminology in the normalization literature) addition takes place after transforming to the proportions and scaling. The scaling factor is usually defined as 10,000 (also used in Seurat by default):

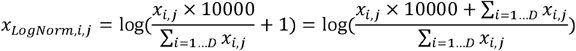 Therefore, for CoDA CLR implementation here, a value of <u>S/10000</u> (S=sum of the counts per cell) was used as the ‘count’ added to the raw counts of each cell (each column):

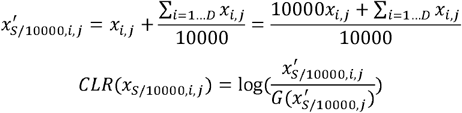
c. Count addition by a fraction with both variable numerator and denominator. Then, we proposed a novel category of count addition. The scaling factor of 10,000 was selected for optimal performance with scRNA-seq data matrix of the current experimental setup of sequencing depth. With increasing sequencing power, the prevailing sequencing depth will increase for future scRNA-seq experiments, therefore both numerator and the denominator should be experiment or results dependent. Thus, we developed the following two count addition methods characterized by having both variable numerator and denominator:
  1. <u>S</u>_j_<u>/Max(S)</u> (S=sum of the counts per cell): Add S_j_/Max(S) to per cell j (every column) raw count, and has a maximum addition of 1.

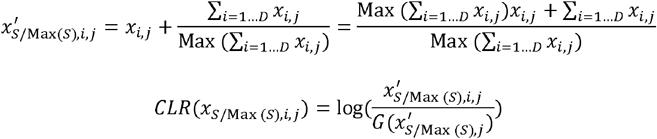
  2. <u>S_j_/GM(S)</u> (S=sum of the counts per cell, GM=geometric mean of per cell (every column) total counts across cells):

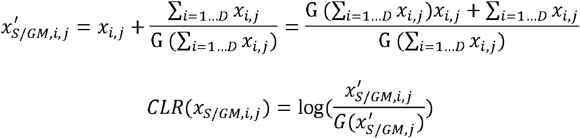 Where *G* (∑ _*i*=1…*D*_ *x*_*i*,*j*_ is the geometric mean of per cell (every column) total counts across cells and 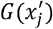 the geometric mean of all genes of cell *j*.

#### 2. Transferability of log-normalized data to CoDA LR transformations

As many datasets have been pre-normalized by conventional log-normalization with scaling, we explored and showed here that such normalizations could be transferable to CoDA by various LR transformations. We introduced an approach to implement count fraction addition with a constant denominator of 10,000 (method b above). CoDA LR transformations based on log-normalization (CLR (LogNorm), which could be served as an approximation) was conducted in log-space, and the main point is that the count of 1 was added to the ‘normalized’ count (proportions scaled to 10,000), followed by logarithm. Specifically for CLR, data was first log-normalized and then subtracted by the arithmetic means of ‘selected genes’ (all genes for CLR):

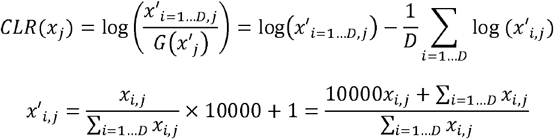

Note: term 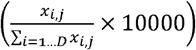 will not affect log-ratios as they are scale invariant.

Then

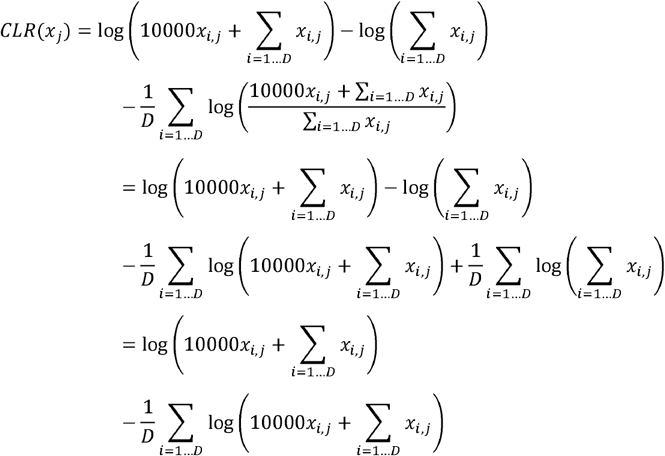

Where *x*_*i*,*j*_ is the raw count of the *i* gene and *j* cell. Term ∑ _*i*=1…*D*_ *x*_*i*,*j*_ served as the ‘count addition’ that differs among cells.

Note that the above step is a special case for sparse scRNA-seq data when performing CoDA and is equivalent as centering the data within each cell (every column) by selected features (e.g., all features for CLR) in log-space. Therefore, the inflated-zero-effect will be mitigated as well. Theoretically, log-ratios in any CoDA data could still be zero when adding a count of 1 when taking the logarithm of zero. However, this is nearly impossible in scRNA-seq data since most of the genes contains zeros across cells and the calculation of the geometric means requires no zeros. It is also worth noting that CoDA of LogNorm with scaling matrix is equivalent to CoDA of S/10000 matrix.

#### 3. Various LR transformations of CoDA evaluated

After various scheme of management of zero counts, the data matrix is ready for CoDA using various kinds of LRs (e.g., CLR, IQLR, HKGLR and ILR) with various advantages and limitations. CLR centered the data by geometric means, while IQLR uses quartile values. HKGLR uses housekeeping gene expressions as reference. ILR is the most robust in CoDA with invariant and isometric properties. Various LRs were then subjected to dimension reduction, clustering and other scRNA-seq downstream tasks for evaluations and were also compared with results generated from typical Seurat workflow using LogNorm or SC-Transform (**Figure 2**). The details of the various LR formulations were shown in **Table S2, Group CoDA**. Note that various LRs use different formulation for the denominator which will affect the output and sometimes makes it difficult for direct interpretation (e.g., ILR)^10^.

1. The most common LR expression of CoDA is the Centered-Log-Ratio (CLR) which takes the geometric mean of all features (genes) as denominators, defined as:

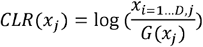 Where *G*(*x*_*j*_)is the geometric mean of the D features (genes) of sample/cell *x*_*j*_.
2. A more complicated LR transformation, Isometric-Log-Ratio (ILR), is defined as:

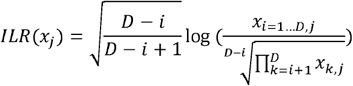
3. A knowledge-based transformation Housekeeping-Gene-Log-Ratio (HKGLR): takes the geometric mean of five housekeeping genes (SDHA, ACTB, UBC, YWHAZ, GAPDH; or other reasonable genes) as the denominator:

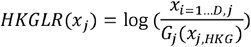

Usually, housekeeping genes have stable expression.

Notably, in addition to the above CoDA LR transformations, some other LR concepts are also well described in the CoDA field or have been proposed previously^7,15,19^, e.g., IQLR, LVHA, mdCLR or group-based LR. We evaluated these transformations as well (data not shown) but found no improvement or worse performance than the most representative transformation CLR, and thus, only the well-known CLR, ILR, and HKGLR were included in the result part for simplicity.

#### 4. Imputation for zero counts

Two algorithms (i.e., MAGIC & ALRA) for imputation of zero counts in scRNA-seq data matrix (**Table S2, Group B**) were also evaluated and compared with CoDA results. Their outputs were CLR-transformed (based on LogNorm) for downstream evaluations as well.

### Data collection and pre-processing

Datasets used in this study were collected from Gene Expression Omnibus (GEO) and other public resources as described in each study^24–39^. Most of these datasets had been used for various algorithm evaluations or benchmark work and were believed to be of high quality^24,29,39,40^. Simulated scRNA-seq datasets were generated by two well-established R packages Splatter^41^ and SplatPop^42^. Details of these datasets were shown in **Table S3**. Processed datasets were collected from previous studies. For the four simulated datasets, we also investigated whether CoDA LR transformations could better reflect the ground truth. We additionally evaluated the performances of various normalizations / LR transformations on dimension reduction and clustering using the ground truth (no dropout) simulated datasets. For raw datasets without prior quality control, we applied the following filtering criteria to remove: (1) Cells expressed < 200 genes; (2) outlier cells based on number of genes expressed (nFeature_RNA), number of total counts (nCount_RNA), and percentage of mitochondrial gene expression (percent.mt); (3) Genes expressed in less than 10 cells.

### Evaluation methods: overview

We evaluated the performance of different normalizations and CoDA LR transformations across several downstream tasks. LogNorm with scaling is the most popular normalization method used in scRNA-seq, so it is used as the baseline for most of the comparison performed here. That is, to compare the performance metrics among different normalizations/LR transformations, we subtracted the metric values by the results of the LogNorm and the difference in performance is shown with box plots, bar plots and heatmaps. Note that some extreme values have been limited to a cutoff value for better visualization in some analyses.

### Dimension reduction and clustering

#### 1. Using partial SVD to replace LRA (easyCODA) in performing CoDA dimension reduction and clustering

CoDA enables unbiased dimension reduction by principal component analysis (PCA) and biplots analysis on log-ratio data^7^. Specifically, the log-ratio analysis (LRA; implemented in easyCODA R Package) is a special case of PCA that applied to the CLR. However, it is not designed for high-dimension data like scRNA-seq matrix, and thus it took a very long time (virtually impractical) to run in ‘easyCODA’. Here, we showed that our CoDA-hd ‘CLR+partial SVD’ process using fast truncated Singular Value Decomposition (SVD) could resulted in an approximation solution--the resulted PCA had similar patterns as those generated by typical CoDA procedure (LRA function) by the ‘easyCODA’ package (**Figure 3** and **Figure S2**). The partial SVD was conducted using the package ‘irlba’, which applies implicitly-restarted Lanczos methods for fast truncated SVD of sparse and dense matrices^43^. Therefore, it provides much better scalability than LRA in ‘easyCODA’.

**Figure 3.**
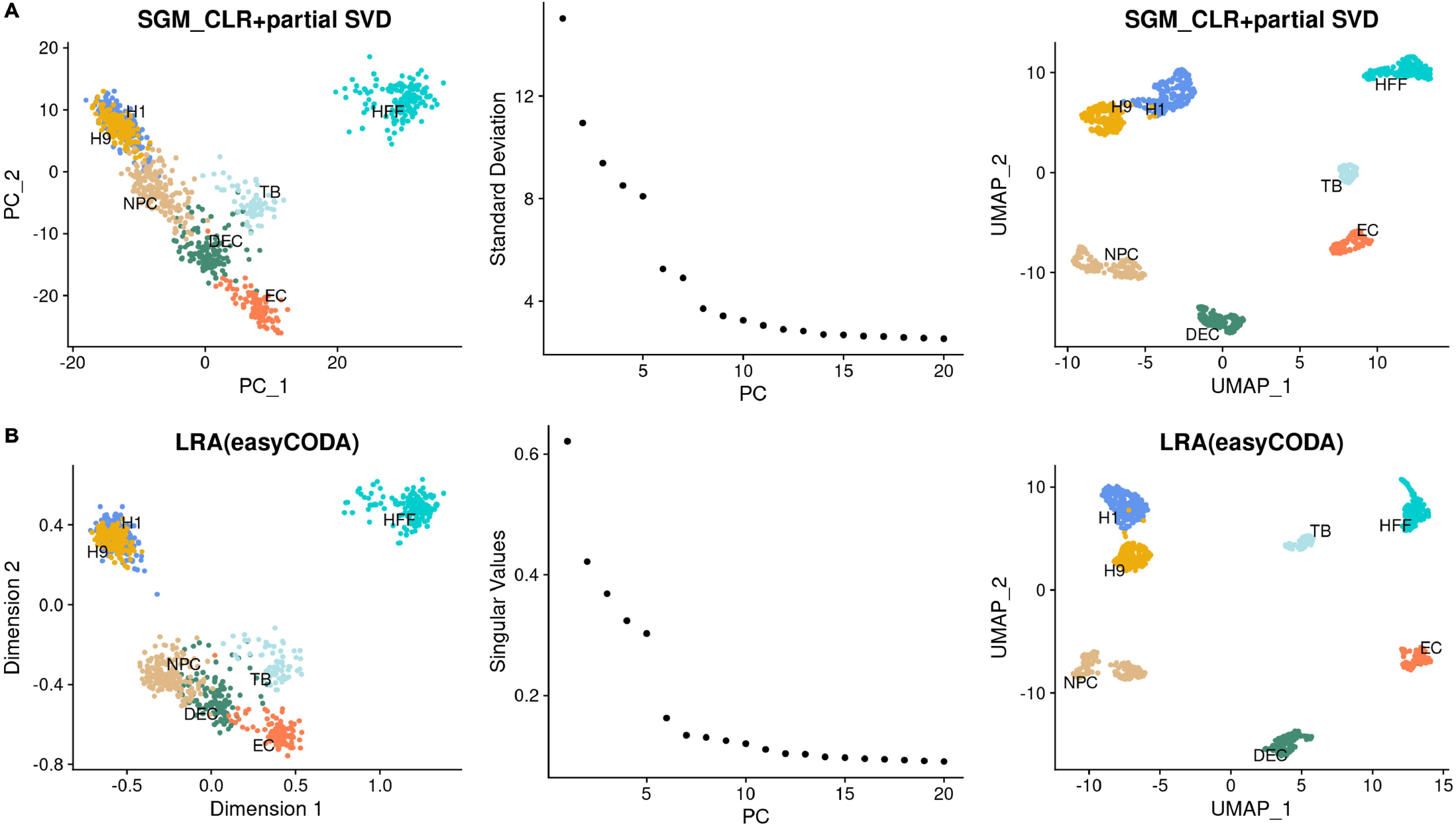
Comparison of the first 2-D PCA & UMAP plots and elbow plots of two dimension reduction algorithms in CoDA--The new CLR+partial SVD vs LRA (typical CoDA dimension reduction algorithm in the easyCODA R-package) using the GSE75748-CellType dataset (top 3,000 features 997 cells). To speed up the dimension reduction process for the high dimensional scRNA-seq data, we perform partial SVD on the CLR (SGM) transformed data and compare the resulted PCs with the standard dimension reduction performed by the LRA command in the easyCODA R package for CoDA data. The first 2-D PCA & UMAP plots and the elbow plots for the dataset GSE75748-CellType for each of the methods are shown for comparison. Overall, the clusters of results are similar between the two algorithms. Therefore, the fast CLR+partial SVD is used in subsequent analysis. (**A**) Results of the partial SVD on CLR (SGM) transformed data. (**B**) Results of the LRA (easyCODA) based on CLR (SGM).

#### 2. Quantitatively evaluate the clustering performance of different normalizations and CoDA LR transformations

We adopted four evaluation metrics, Entropy of accuracy (*H*_*acc*_), Entropy of purity(*H*_*pur*_), Adjusted Rand Index (ARI) and Normalized mutual information (NMI)^29,40^ with K-means Clustering and Louvain Clustering algorithms on multiple labelled (i.e., with ground truth cell type label) datasets. For clustering, the number of clusters was set to be the number of known cell type labels in each dataset. The top 3,000 variable genes and the top 10 PCs were used for the cell clustering. The detailed implementations of the clustering algorithms were described previously^29^.

#### 3. Metrics to access clustering concordance with the ground truth

*H*_*acc*_ evaluates the difference of the true groups within each predicted cluster, defined as:

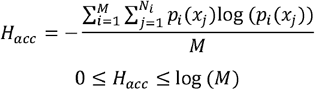

where *M* is the number of predicted clusters; *N*_*i*_ is the number of true groups in the *i*^*th*^ predicted cluster; *x*_*i*_ are cells in the *i*^*th*^ true group; and *p*_*i*_(*x*_*j*_) are the proportions of cells in the *i*^*th*^ true group relative to the total number of cells in the *i*^*th*^ predicted cluster.

_*Hpur*_ evaluates the difference of the predicted clusters within each true group, defined as:

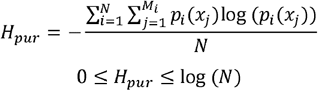

where *N* is the number of true group; *M*_*i*_ is the number of predicted clusters in the *i*^*th*^ true group; *x*_*i*_ are cells in the *i*^*th*^ predicted cluster; and *p*_*i*_(*x*_*j*_) are the proportions of cells in the *i*^*th*^ predicted cluster relative to the total number of cells in the *i*^*th*^ true group. Smaller *H*_*acc*_ and *H*_*pur*_ indicates better clustering performance^29^.

Adjusted Rand index (ARI) that evaluates the similarities between two data distributions^40^ was calculated using adjustedRandIndex() function in mclust package^44^, defined as:

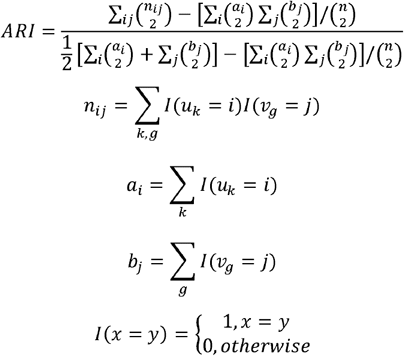

Where *i* and *j* enumerate the *k* clusters; 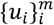 is the predicted cluster label; 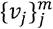 is the true group label.

Normalized mutual information (NMI) that measures the correlation between two random variables^40^ is calculated using NMI() function from aricode package^45^, defined as:

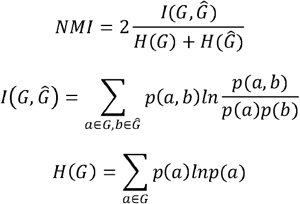

Where *G* is the true group, 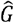 is the predicted cluster.*p*(*a*), *p*(*b*) and *p*(*a*,*b*) are the probabilities that the cell belongs to cluster a, cluster b and both, respectively. For both ARI and NMI, higher values indicate better clustering performance.

To compare the above metrics among different methods, we first subtracted all the ARI and NMI values by the results of the log-normalized data (vice versa for *H*_*acc*_ and *H*_*pur*_ for better visualization) and then took the median values of the four real or simulated datasets. The heatmaps were generated for both median aggregates and individual dataset (note that the extreme values have been limited to a specific cutoff for better visualization). Next, 2-D PCA or UMAP plots were generated for each dataset to evaluate the performance of different methods on dimension reductions. The first 10 PCs were used to generate the UMAP.

For the four simulated datasets, we further evaluated the performances of various normalizations / LR transformations using the ground truth (no dropout) simulated datasets.

### Effect of Zero-inflation on dimension reduction

Low-quality cells (e.g., degraded cells, cell debris) are common in scRNA-seq datasets. To simulate the effect due to the presence of degraded cells, we experimented with spike-in 10% of zero-inflated cells into high-quality datasets CellBench-10X-5CL and GSE75748. For example, 10% of the H1975 cells (a cell line originated from a lung tumor) were randomly selected and copied to simulate low-quality cells. For each copied cell, 60% of the genes (∼6000 genes) were randomly assigned to zero. Thus, the new dataset has 10% more degraded H1975 cells with at least 70% features set to zero (i.e., zero-inflated vectors). The data was then processed by various normalizations and CoDA LR transformations, followed by the evaluations on PCA and UMAP. Additionally, their clustering performance were evaluated based on the subset cell of H1975 and H1975-zero. Similarly, 40% of the H1 cells from dataset GSE75748-CellType were copied and simulated as described above. The H1 and H9 cell subset was then evaluated.

### Trajectory inference

To evaluate the performance of different methods on trajectory inference, we applied Slingshot^46^, DPT^47^, Monocle2^48^ and Monocle3^49^ on twenty-two real datasets with known cell-state/time labels (**Table S3**). Spearman correlation coefficients (SCC) were calculated between the predicted pseudotime and the ground truth time labels. Another metric Pseudo-temporal ordering score (POS) which measures cell orders were calculated using orderscore() in TSCAN^50^ package. Due to space limitation and scalability, fifteen ‘gold standard’ datasets from Saelens et al.^39^ were not used in MAGIC and ALRA imputation.

In Slingshot, the starting point of the trajectory was set based on the known starting time and other parameters were set to default. For trajectory with multiple branches, the predicted branches and the real branches were matched by their largest overlaps of cells. Specifically, for datasets with three cell types (A, B, and C where A will differentiate to B and C) with even number, ideally predicted branch 1 would have 100% overlap rate with A-to-B known branch while only have 50% overlap rate with A-to-C known branch (vice versa for predicted branch 2). Then the true overlap rates (i.e., largest overlaps between predicted branches and real branches, as matched branches), trajectory false rates (average of percentage of cells in selected real branch that appear in other predicted branches), incorrect cell type proportions (percentage of cells in matched predicted branch that did not belong to matched real branch) were calculated. 2-D PCA plots were generated for visualizing Slingshot results.

In DPT, same procedures were applied and the predicted branches by DPT were further evaluated with clustering metrics *H*_*acc*_, *H*_*pur*_ ARI, and NMI (as described in Clustering). 2-D diffusion map plots were generated for visualizing trajectories. In Monocle2 & 3, same procedures were applied.

### Biomarker predictive performance

Ratio-based biomarkers have been reported to have great performance in classifications or predicting disease status^51,52^. Indeed, researchers have used the ratios of gene abundances in qPCR data to study the expression differences between conditions. In this study, we evaluated the performance of different transformations in cell type biomarker identification using three real scRNA-seq datasets with matched bulk (GSE75748, GSE81861, CellBench-10X-5CL) and the three simulated datasets 1-3 (**Table S3**). Top 10 differential expressed genes (DEGs) between different cell types identified in bulk data (i.e., totally 410 marker genes) or ground truth DEGs in simulated datasets were used for evaluation in scRNA-seq data. The predictive performance was evaluated by calculating the area under the receiver operating characteristic curve (AUROC) for each of the selected genes in each comparison. We determined to use AUC here as it is a better metric that does not have any statistical assumption, while other differential expression analysis methods have different statistical basis / assumptions which may not be applicable for compositional data or other types of data. The AUC values were then compared among different normalizations / LR transformations. Additionally, the percentages of marker genes that had AUC greater certain cutoffs were calculated for comparison. In simulated datasets, 50 known non-marker genes (ground-truth non-DEGs) between pairwise cell types were randomly selected out and their AUC were calculated for studying potential false positive rates.

### Other statistical analysis

For each of the comparisons with enough data points, Wilcoxon rank sum tests were performed between the results of the selected methods and the conventional log-normalization. The significance levels were shown in each box plot with *, **, *** represents ‘P<0.05’, ‘P<0.01’ and ‘P<0.005’ respectively, and ‘#’ stands for not-significant-difference (‘P<0.1’).

### Running time and memory usage

We used four datasets (10000 × 1500, 10000 × 5000, 10000 × 10000 and 10000 × 50000 matrix) to evaluate the time spent and memory usage of different methods (Log-normalization, SC-Transform and CLR). Additionally, we compared the running time and memory usage between LRA-easyCODA and CLR-partial-SVD. The top 3,000 variable features were used in LRA and partial-SVD. To obtain the maximum memory usage, we used the peakRAM() function from the ‘peakRAM’ package.

## Results

We first evaluated different count addition methods on the performance of cell clustering and projection after dimension reduction. By default, various methods were compared against log-normalization, which was used as the baseline for comparison. Thus, difference in performance in other methods are shown as gain or loss over the log-normalization, the default method commonly used.

### CoDA LR-based transformations improve clustering performance

Quantitative evaluations of clustering were performed using four real datasets and four simulated datasets for comparison among various normalizations and LR transformations. **Figure 4** evaluated the effect of applying CLR transformation to various ‘count addition’ schemes, while **Figure 5** compared the performance of various LR transformations of CoDA, namely CLR, ILR and HKGLR. CoDA LR transformations, particularly CLR using SGM count addition, showed significant improvements in all clustering metrics (H_acc_, H_pur_, ARI, NMI) compared to conventional normalizations using either Louvain or K-means algorithms for both real and simulated datasets (**Figure 4-5** and **Figure S3-4**). CLR (LogNorm / S10000) also performed well in these evaluations. In contrast, constant count addition schemes (i.e., P1-CLR, P0.01-CLR, P0.0001-CLR) performed substantially worse. This confirms our hypothesis that cell-specific count addition based on total counts is more effective than simple constant addition and is also a prerequisite for successful CoDA-hd. Importantly, we demonstrated that log-normalized data could be directly transformed to approximate CoDA LR representation through CLR (labelled as LogNorm-CLR in figures). This provides a straightforward way to apply CoDA to pre-normalized datasets by LogNorm that are commonly available in public repositories. Therefore, the benefit of CoDA-hd is readily applicable to most public datasets.

**Figure 4.**
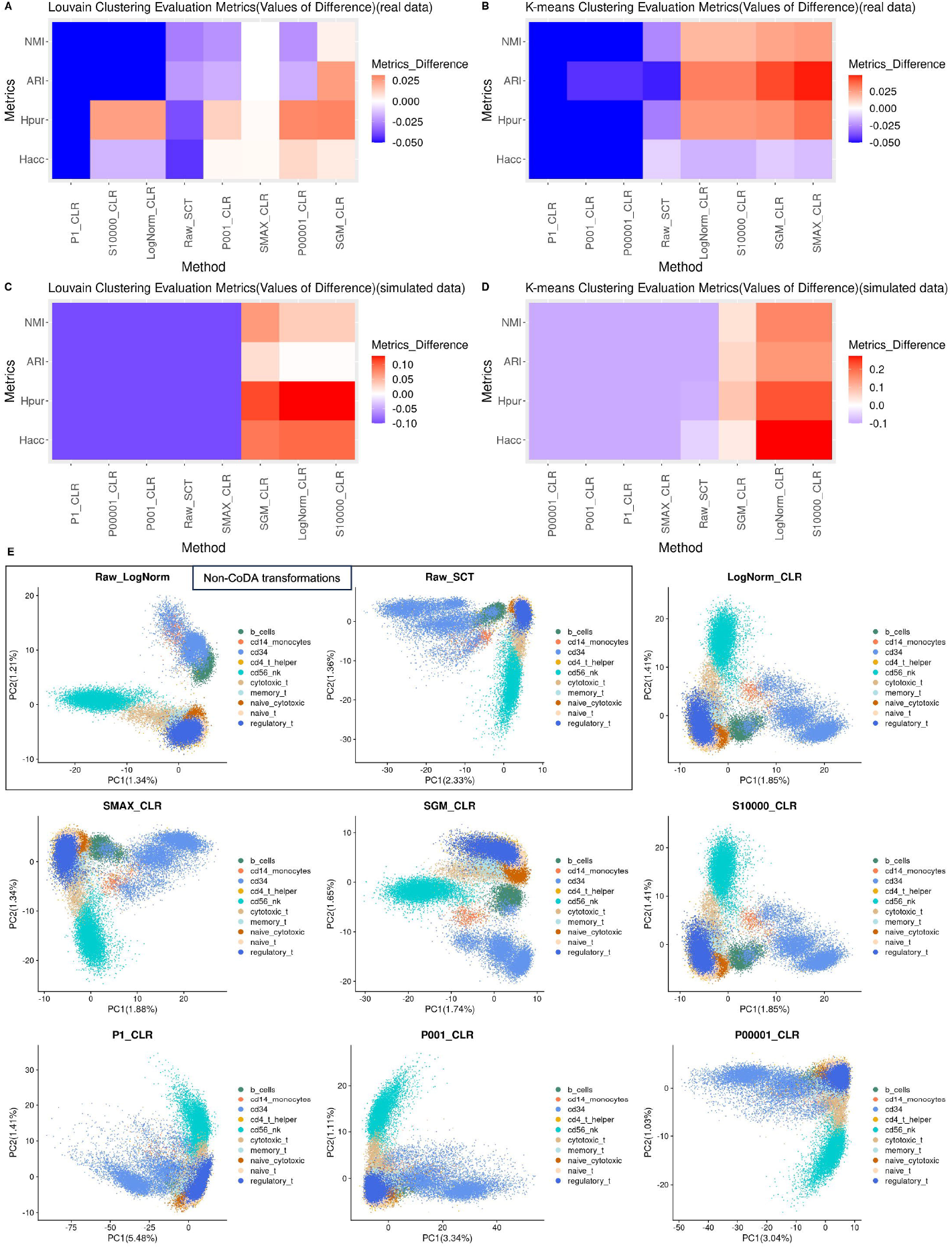
Handling of zero counts: clustering performance of different count addition schemes in CoDA transformations using K-means and Louvain Clustering algorithms on real and simulated datasets and the first 2-D PCA plots using the sorted PBMC dataset. Four evaluation metrics of clustering performance, Entropy of accuracy (*H*_*acc*_), Entropy of purity (*H*_*pur*_), Adjusted Rand Index (ARI) and Normalized mutual_pur_ information (NMI) are used to quantitatively evaluate the clustering performance of different CoDA count addition schemes by Louvain Clustering and K-means Clustering. Four datasets with given cell-type labels (ground-truth) are used. The number of clusters is set to the number of known cell type labels in each dataset. Top 3000 high variable genes are used for running PCA. The top 10 PCs are used for clustering. ARI and NMI are subtracted by the results of the raw log-normalization (as baseline) (vice versa for *H*_*acc*_ and *H*_*pur*_ for better visualization) and the medians are taken. Extreme values in heatmaps have been limited to a cutoff for better visualization. CLR count addition schemes include: P1: plus 1; P001: plus 0.01; P00001: plus 0.0001; SMAX: plus Sum(j)/Max(Sum(j)); SGM: plus Sum(j)/Geometric-Mean(Sum(j)); S10000: plus Sum(j)/10,000; LogNorm_CLR: CLR transformed from log-normalized data. CLR transformation of log-normalized data (logNorm_CLR) produces better clustering than log-norm itself (Raw-LogNorm). Overall, those schemes adding a constant value (P1, P001, P00001) result in cell clusters that are less well demarcated. (**A-B**) Median metric difference values of the four real datasets using K-means and Louvain Clustering algorithms. (**C-D**) Median metric difference values of the four simulated datasets using K-means and Louvain Clustering algorithms. (**E**) First 2-D PCA plots of different CoDA count additions using the cell type sorted PBMC dataset.

**Figure 5.**
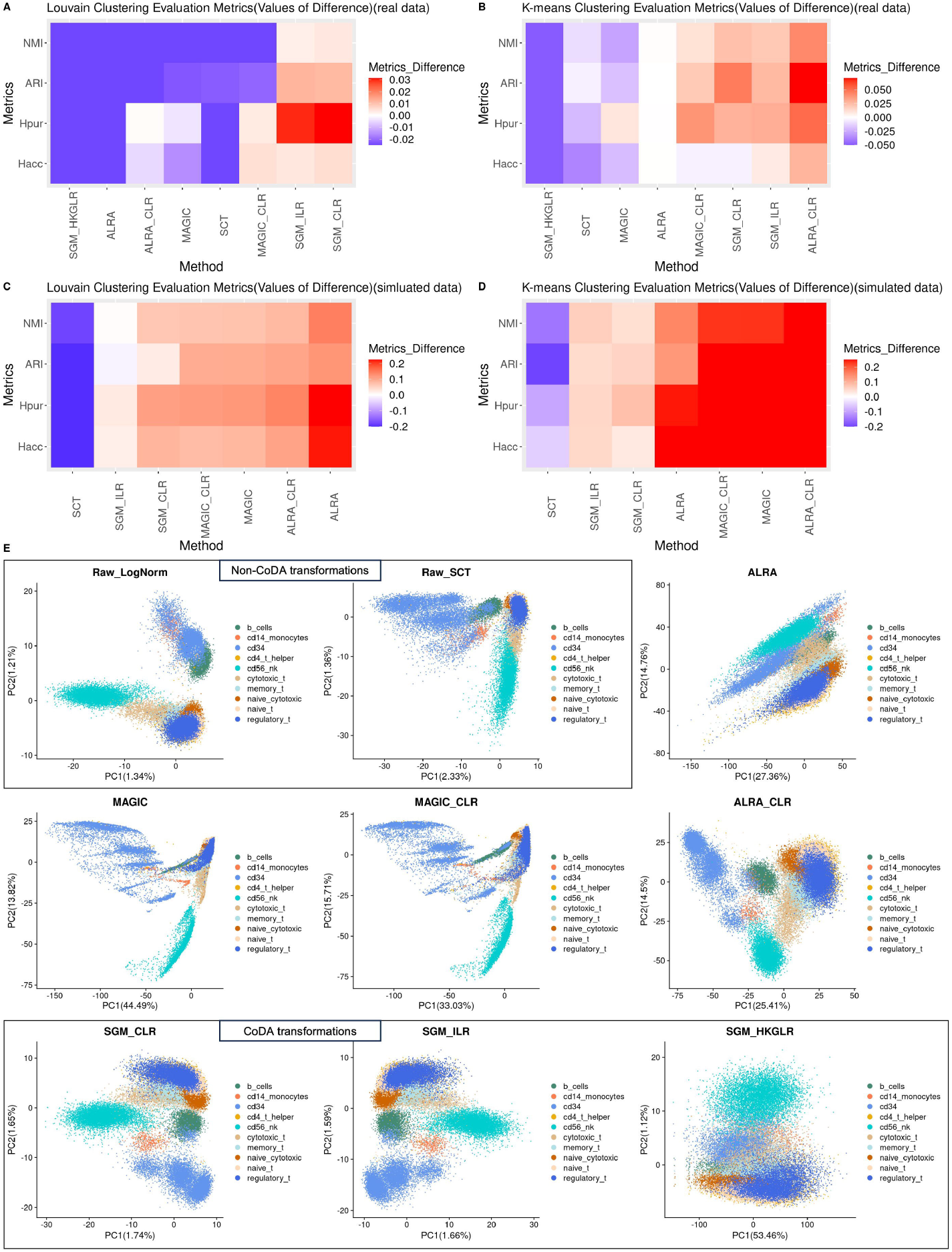
Clustering performance of different transformations / imputation algorithms with or without CoDA transformations using K-means and Louvain Clustering algorithms on real and simulated datasets and the first 2-D PCA plots using the sorted PBMC dataset. Four evaluation metrics of clustering performance, Entropy of accuracy (*H*_*acc*_), Entropy of purity (*H*_*pur*_), Adjusted Rand Index (ARI) and Normalized mutual information (NMI) are used to quantitatively evaluate the clustering performance of different normalizations / CoDA transformations / imputation algorithms by Louvain Clustering and K-means Clustering. Four datasets with given cell-type labels (ground-truth) are used. The number of clusters is set to the number of known cell type labels in each dataset. Top 3,000 high variable genes are used for running PCA. The top 10 PCs are used for clustering. ARI and NMI are subtracted by the results of the raw log-normalization (as baseline) (vice versa for *H*_*acc*_ and Entropy of purity (*H*_*pur*_), Adjusted Rand Index (ARI) and Normalized mutual for better visualization) and the medians are taken. Extreme values in heatmaps have been limited to a cutoff for better visualization. SGM_CLR represents the CLR transformation with count addition ‘Sum(j)/Geometric-Mean(Sum(j))’. Imputation algorithms produce extra spurious correlation between PC1 and PC2 (e.g., ALRA, MAGIC) which are not found in the datasets. Overall, CLR transformation of imputed data shows better clustering except in the simulated dataset undergone Louvain clustering. (**A-B**) Median metric difference values of the four real datasets using K-means and Louvain Clustering algorithms. (**C-D**) Median metric difference values of the four simulated datasets using K-means and Louvain Clustering algorithms. (**E**) First 2-D PCA plots of different normalizations / CoDA transformations / imputations using the sorted PBMC dataset.

On the other hand, imputation by ALRA or MAGIC alone (labelled as MAGIC and ALRA in **Figure 5**) did not perform well for real datasets. However, the CoDA LR transformations of these imputed matrices showed great improvement (comparing MAGIC-CLR vs MAGIC or ALRA-CLR vs ALRA). This suggests that the CoDA framework enhances the utility of imputation approaches.

Using simulated datasets, we also compared the metrics among various normalizations and count-addition CLRs against their ground truth (no dropout) datasets (i.e., True-LogNorm and True-CLR). As expected, True-SGM-CLR and True-LogNorm-CLR consistently outperformed True-LogNorm (**Figure S5**). Therefore, we found that SGM-CLR was an optimal count addition scheme and it was used in other downstream analyses.

### CoDA LR-based transformations improve projections and visualizations of scRNA-seq datasets with known cell-type identities

In PCA and UMAP visualizations, CLR (SGM) and CLR (LogNorm / S10000) resulted in more discrete and biologically coherent cell-type clusters (**Figure 4** and **Figure S6**). Compared to LogNorm alone, cell-types, especially monocytes, were better separated using CLR (**Figure 5**). On the other hand, knowledge-based denominator selection (HKGLR) performed poorly, creating complex and potentially misleading structures. An important observation provided by CoDA was that certain cell-type relationships appeared more biologically plausible with CLR. For example, in UMAP projection of the PBMC data, naïve cytotoxic cells appeared far from cytotoxic cells when using log-normalization, while with CLR the two related cell types were appropriately connected, reflecting their biological relationship (**Figure S7**).

When comparing the two zero-imputation methods, MAGIC and ALRA resulted in spurious, complex and sharp cell-type structures in PCA which were not apparent in log-normalization or CLR. Interestingly, CoDA LR transformations of ALRA provided better projections of cell-types especially in PCA (**Figure 5E**).

Similarly, the cell-types in CellBench-10X-5CL dataset shown in **Figure S8-11** can be easily clustered and projected in PCA/UMAP by most of the normalizations / CoDA LR transformations. CLR (SGM) consistently performed well in these projection visualizations. In UMAP, H1975 cell lines may be divided into subclones as they appear in separated clusters in some projections (**Figure S11**). Therefore, it is selected for zero-inflation (degraded cells) spike-in experiment. Again, MAGIC generated most farther separated sub-clusters of the H1975 cell line which may represent an artefact due to imputation.

In simulated dataset 2, we included the log-normalization and CLR of ground truth dataset (no dropout) as baseline (i.e., True-LogNorm and True-CLR). Interestingly, in 2-D PCA, though cells were mixed with each other and no obvious pattern was observed for most of the methods (even for ground truth log-normalization), we observed better separations of clusters in True-CLR and ALRA-CLR, compared to their log-normalized formulation (**Figure S12**). This again highlighted the advantage of CLR over the log-normalization. In 2-D UMAP, True-LogNorm and True-CLR produced distinct and concentrated cell-type clusters (**Figure S13**). For the dropout dataset, CLR resulted in more distinct and concentrated clusters compared to log-normalization and were more similar to the ground truth.

### CoDA provides better clustering for degraded cells

Since low-quality cells with high dropout rates may cause misleading conclusions, we next performed a simulation analysis in which we spiked in degraded cells (10% of copied H1975 cells with zero-inflated parts) in the high-quality dataset CellBench-10X-5CL (see Methods). We studied whether CoDA LR transformations could provide better clustering of these degraded zero-inflated cells which is essential to avoid erroneous results. In both 2-D PCA and UMAP visualizations, H1975-copied-zero cells (orange dots in **Figure 6A** and **Figure S14A**) were separated from original H1975 cells to a different extent by various methods except SC-Transform (**Figure 6A**). However, only CLR and ILR can distinctly cluster these artificially degraded cells which is a desirable feature. On the other hand, log-normalization presented these degraded cells as scattered between distinct cell-type clusters, which could easily lead to misinterpretation of these degraded cells as transitional states between clusters. In contrast, CLR clearly identified these as a separate cluster of low-quality cells. This finding is particularly important as it suggests that CoDA may help prevent the misidentification of technical artifacts as biological transitions. It is indicated that low-quality cells with high dropout rate are possible to raise false trajectories between different clusters, and CoDA may be a better alternative than conventional log-normalization in handling scRNA-seq data. In HKGLR UMAP, the degraded cells were spuriously split into two clusters and all cells were heavily scattered, which was repeatedly observed in different spike-in runs. Therefore, the use of knowledge-based denominator selection for CoDA transformations of scRNA-seq is not supported.

**Figure 6.**
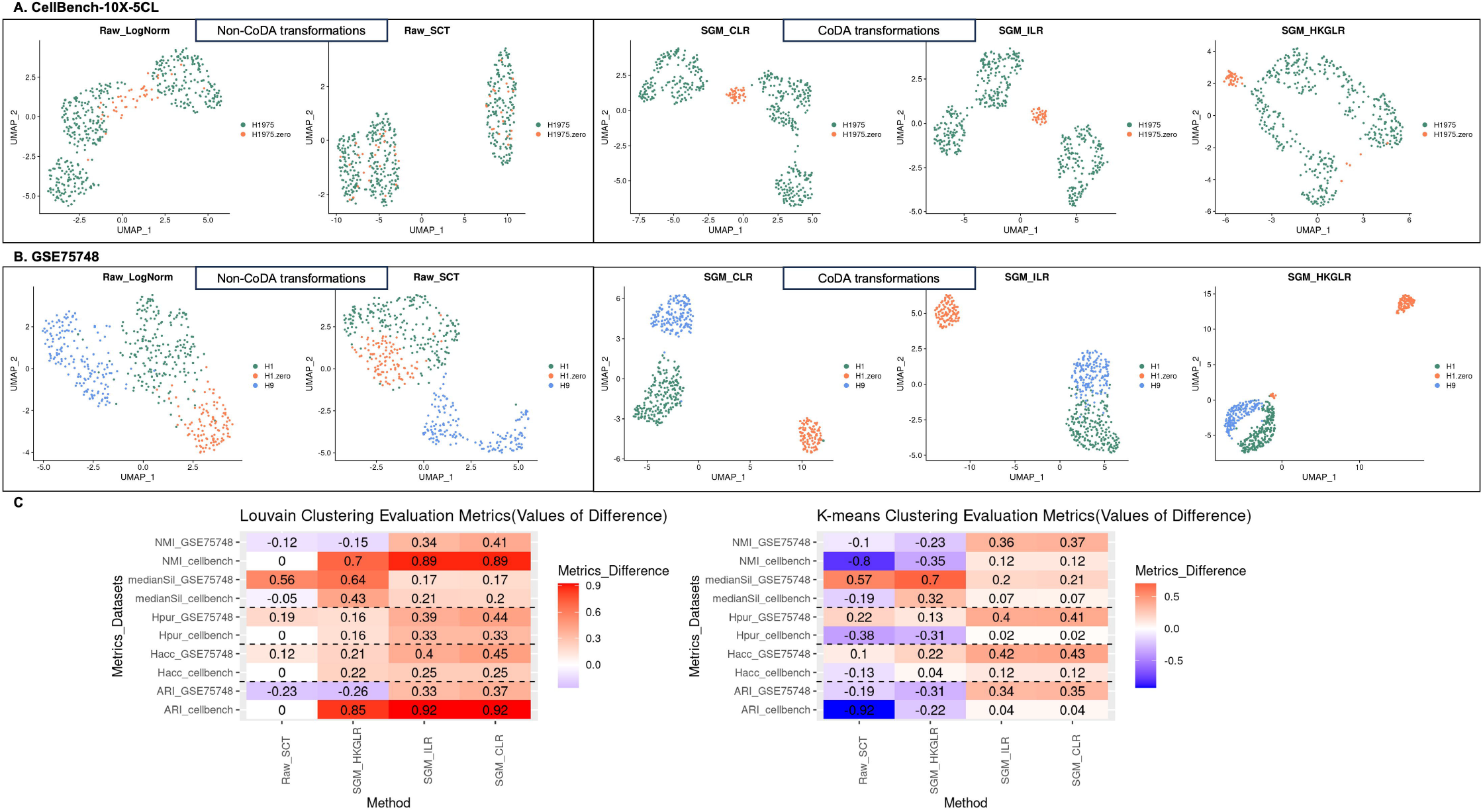
Effect of degraded cells: performances of different normalization methods and CoDA LR transformations when degraded cells are present. **(A)** 2-D UMAP plots of the CellBench-10X-5CL H1975 and H1975-zero subset processed by different normalizations and CoDA transformations. For the dataset CellBench-10X-5CL, 10% of the H1975 cells are randomly selected and copied to simulate low-quality cells. For each copied cell, 60% of the genes (∼6000 genes) are randomly assigned to zero and then spike-in the data matrix as additional cells. Thus, the new data matrix has 10% more degraded H1975 cells with at least 60% feature (genes) counts set to zero. The data is then processed by various normalizations and CoDA LR transformations, followed by the evaluations on PCA and UMAP. For UMAP visualization, top 10 PCs are used. UMAP plots for H1975 cell subset are generated. Additionally, clustering performances are evaluated based on the separation of H1975 cells and the degraded cells (labeled as H1975-zero in the figure). **(B)** 2-D UMAP plots of the GSE75748-CellType H1, H1-zero, and H9 cell subset. Similarly, 40% of the H1 cells from dataset GSE75748-CellType are copied and simulated as spike-in degraded cells as described above. The separation of H1, H1-zero, and H9 cell clusters are then evaluated. (**C**) Results of the clustering performances of various normalizations / CoDA transformations on the two degraded cell datasets using Louvain and K-means algorithms. ARI and NMI are subtracted by the results of the raw log-normalization (as baseline) (vice versa for *H*_*acc*_ and *H*_*pur*_ for better visualization).

Similarly, in the GSE75748-CellType dataset, CoDA LR transformations better clustered the H1, H9 and degraded H1-copied-zero cells compared to conventional methods (**Figure 6B** and **Figure S14B**).

The quantitative evaluations confirmed that CLR showed improvements in all clustering metrics when applied to these two subset zero-inflated datasets (**Figure 6C**). This robustness to dropout effects represents a major advantage of CoDA for scRNA-seq analysis.

### CoDA improves trajectory inference

When evaluating trajectory inference across twenty-two real ‘gold standard’ datasets with known time labels, CLR showed statistically significant improvements in both Spearman correlation coefficients (SCC) and Pseudo-temporal ordering score (POS) compared to log-normalization using Slingshot (**Figure 7A-B** and **Figure S15**). Interestingly, ALRA-CLR showed substantial improvement compared to ALRA alone.

**Figure 7.**
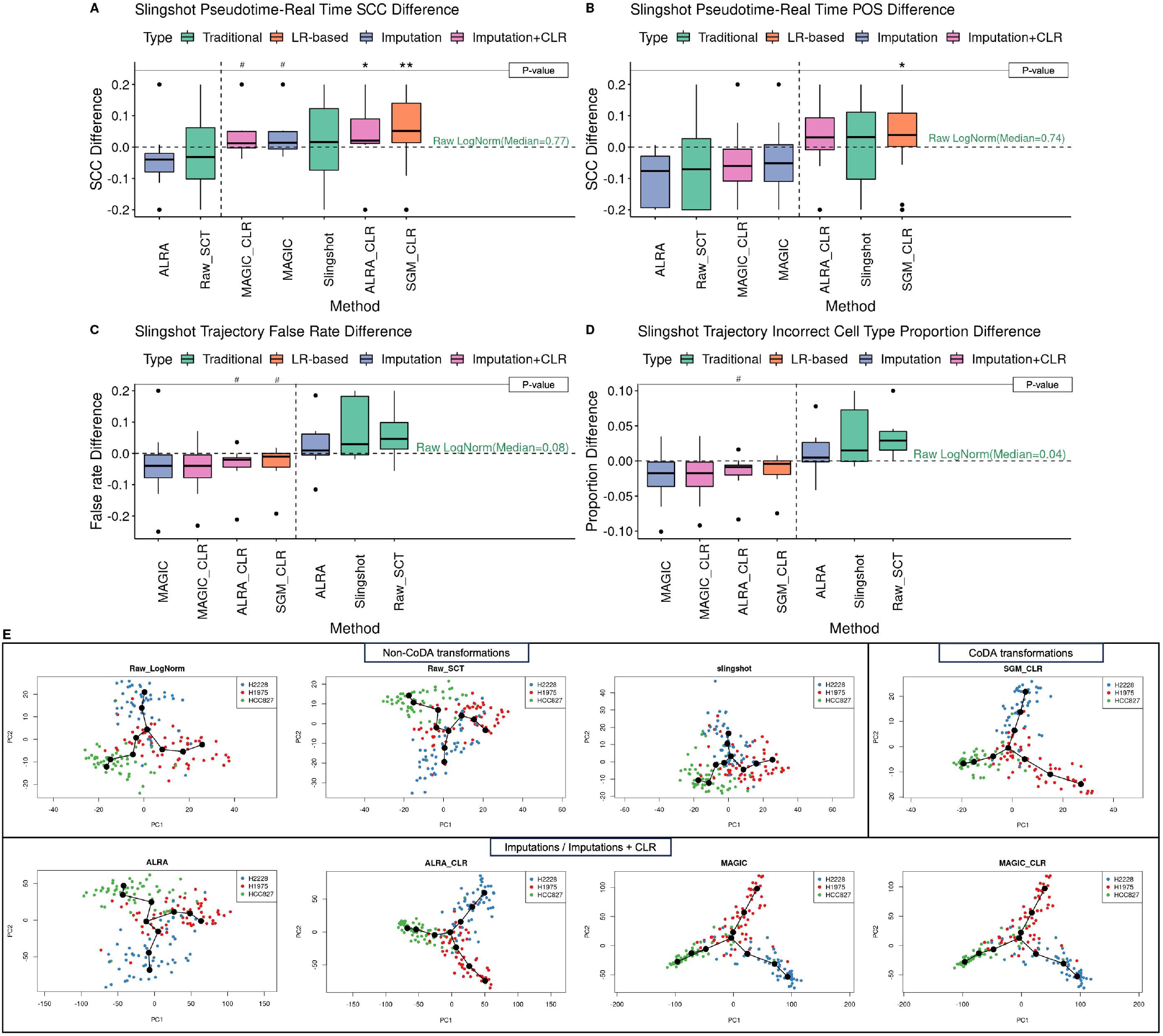
Comparison of effects of various normalizations / CoDA transformations on Slingshot trajectory inference. The performances of different normalizations / imputations / CoDA transformations on trajectory inference are evaluated with Slingshot using twenty-two datasets with real time & lineage (ground-truth) labels. The starting point of the trajectory is set based on known information and other parameters are set to default. For trajectory with multiple branches, the predicted branches and the real branches are matched by their largest overlaps of cells (see Methods). *, ** and *** represent ‘P<0.05’, ‘P<0.01’ and ‘P<0.005’ respectively and ‘#’ stands for not-significant-difference (‘P<0.1’), when compared to the baseline method raw log-normalization using Wilcoxon rank sum test. (**A-B**) Spearman correlation coefficients and Pseudo-temporal ordering scores are calculated between the predicted pseudotime and the real time labels. ALRA-CLR and SGM-CLR show significant improvement over baseline and Slingshot-default-transformation. (**C-D**) Generation of false results analyzed by using the trajectory false rates and the incorrect cell type proportions. Three CLR schemes in addition to MAGIC may have better (not statistically significant) control of false results generation. (**E**) The 2-D Slingshot PCA plots generated for the CellBench-10X cellmix3 dataset after different normalizations / imputations / CoDA transformations to visualize trajectories.

For datasets with multiple branches, CLR had lower false trajectory rates and incorrect cell type proportions compared to log-normalization using Slingshot (**Figure 7C-D** and **Figure S15**). Visualization of trajectories (e.g., CellBench cellmix3, **Figure 7E**) showed that CLR presented more concentrated and coherent trajectories, with cells from the same cell types appearing closer together and forming clearer continuums.

Similar improvements were observed with DPT, where CLR enhanced trajectory inference with statistical significance (**Figure S16**). In Monocle2, CLR consistently performed better than conventional normalizations and imputations (**Figure S17**). However, in Monocle3, CLR showed no improvement but maintained consistent performance with log-normalization.

### Enhanced identification of biomarkers after CoDA LR transformation

In the evaluation of biomarker identification, imputation methods (MAGIC, ALRA and their CLR-transformations) consistently improved true positive rate (**Figure 8**). CLR showed modest improvements when considering all differentially expressed genes (DEGs) (**Figure 8A**), but demonstrated significantly higher AUC values for markers that performed poorly with log-normalization (LogNorm AUC<0.75 (**Figure 8B**) or AUC<0.9 (data not shown)).

**Figure 8.**
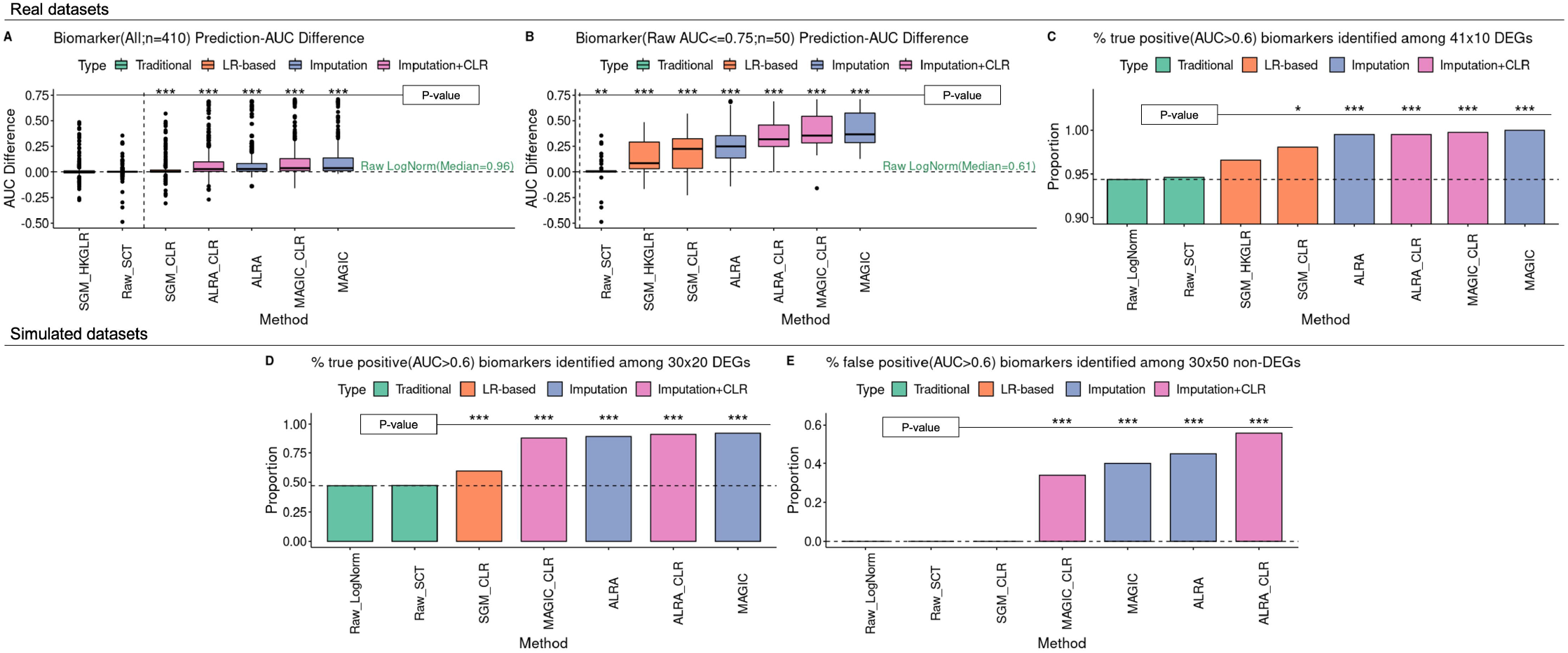
Performances of various normalizations / CoDA transformations on cell-type-specific biomarkers (DEGs) detection. The biomarker detection performances of different normalizations / imputations / CoDA transformations are evaluated. Top 10 DEGs between pairwise cell types in bulk data are used as real biomarkers in three real datasets (**A-B**). *, ** and *** represent ‘P<0.05’, ‘P<0.01’ and ‘P<0.005’ respectively, when compared to the baseline method raw log-normalization using Wilcoxon rank sum test. (**A**) The AUCs for each method for each of the biomarkers are subtracted by the values of the raw log-normalization (baseline). AUCs of the biomarkers improve by CoDA transformations (SGM-CLR, MAGIC-CLR and ALRA-CLR) and imputation methods (ALRA, MAGIC). (**B**) When focusing on those DEGs that have only moderate AUC <=0.75 on log-normalized data matrix, four CoDA transformations (SGM-HKGLR, SGM-CLR, ALRA-CLR, MAGIC-CLR) improve AUCs significantly. Both imputation methods also improve AUCs. (**C-D**) If AUC>0.6 is used as cutoff for useful biomarkers, CoDA and imputation methods may enhance identification of biomarkers in real dataset (C) and simulation dataset (D). (**E**) However, in the simulation dataset with ground-truth, false positive biomarkers are only under control by Raw-LogNorm (as baseline), Raw-SCT and SGM-CLR. 30% to 50% of the selected non-DEGs are identified as potential biomarkers by ALRA or MAGIC (even with CLR transformation).

Both LR transformations and imputations resulted in higher percentages of markers with AUC greater than 0.6 (**Figure 8C-D**), 0.75 or 0.9 (data not shown). This suggests that CoDA can enhance the performance of challenging biomarkers that conventional normalization struggles to identify.

Importantly, while imputation methods also increased the false positive rate (higher AUC for known non-markers (non-DEGs) in simulated dataset), CoDA LR transformations maintained specificity similar to conventional normalization (**Figure 8E**). This balance of improved sensitivity without increased false positives represents another advantage of CoDA.

### Running time and memory usage

As expected, log-normalization was the fastest method regardless of cell number, while SC-Transform was the slowest. CLR required more computational resources than log-normalization but less than SC-Transform (**Figure 9A-B**). For dimension reduction, our ‘CLR+partial-SVD’ approach was substantially faster and more memory-efficient than standard LRA implementations, making CoDA feasible even for large datasets (**Figure 9C-D**). Both procedures completed within hours using up to 35 GB RAM which are typical setup in servers and even on desktop computer. Such simple computing power requirement enable CoDA-hd to be used by most researchers.

**Figure 9.**
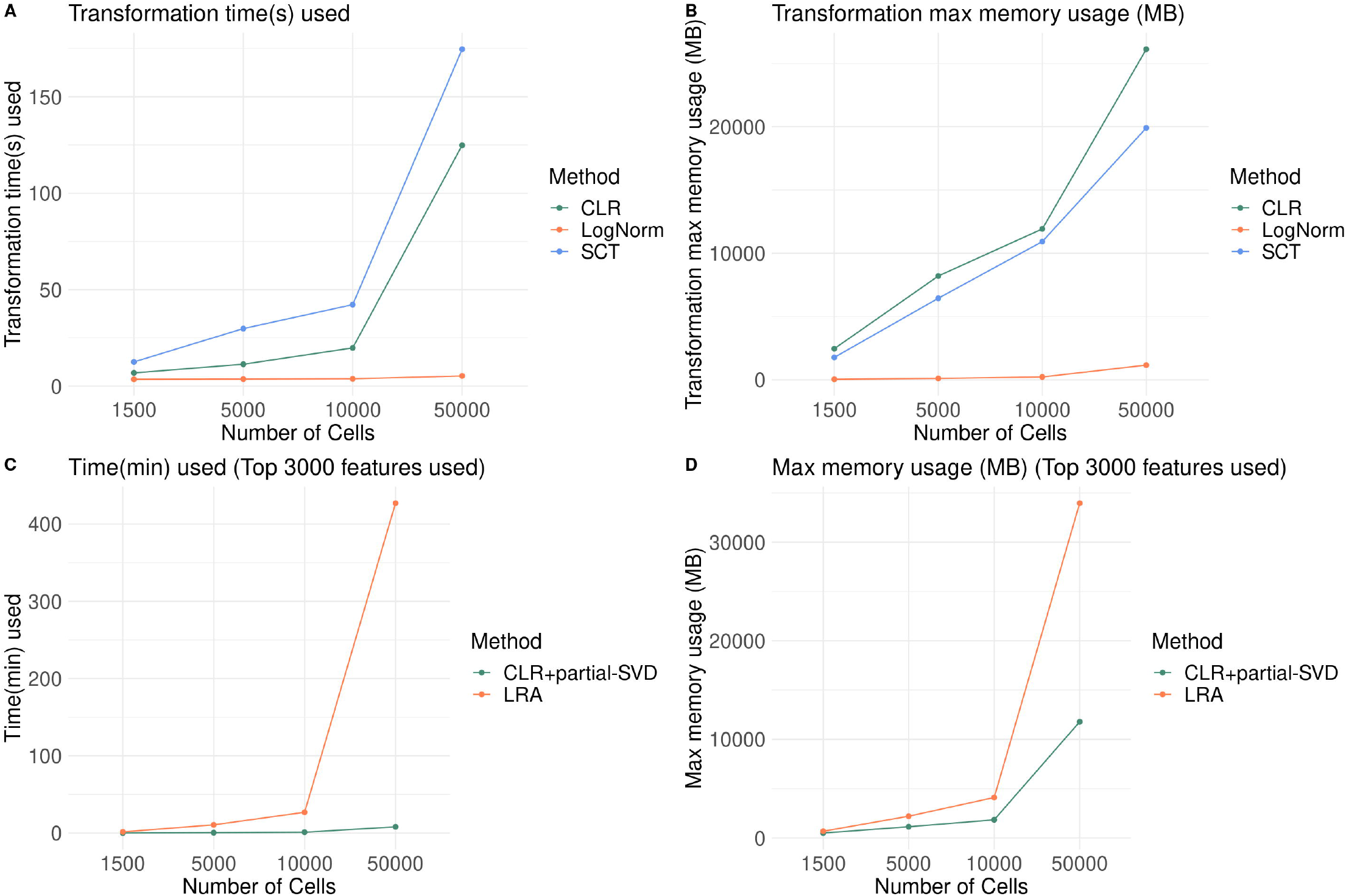
Computer resources (running time and memory requirement) needed for CoDA transformation (CLR) and analysis (CLR-partial-SVD and LRA). (**A**-**B**) Running time (A) and memory requirement (B) for conventional log-normalization, SC-Transform and CLR transformation are compared. All of them are dependent on the number of cells in the data matrix. Four sizes of datasets (10000×1500, 10000×5000, 10000×10000 and 10000×50000 matrix) are applied. For the largest data matrix (10000×50000 matrix), CLR transformation can be completed within minutes using 20 GB RAM. (**C**-**D**) Running time (C) and memory requirement (D) for CLR+partial-SVD and LRA (easyCODA) are compared. Time required (C) and memory usage (D) for performing CLR-partial-SVD are much smaller than conventional LRA in the easyCODA package.

## Discussion

We successfully applied the complete workflow of CoDA from data closure, LR transformation to dimension reduction. These procedures are also shown to be compatible with commonly used downstream analysis algorithms of scRNA-seq data. The two key innovations in our study are (1) SGM count addition scheme developed specifically for scRNA-seq and may be for other sparse data matrix and (2) CoDA of high dimensional data. Using more than thirty real or simulated datasets, we demonstrated some advantages of CoDA LR transformations, especially CLR with SGM count addition, in multiple downstream analyses of scRNA-seq.

We pioneered new schemes of count addition (e.g., SGM count) and CLR after closure of data treated by log-normalization. These two CoDA techniques improved data representation of scRNA-seq data matrix. The use of partial SVD also allows PCA of LR to be finished seamlessly for high dimensional (hd) matrix in a matter of minutes, in contrast to the standard CoDA LRA algorithm which is designed for smaller datasets.

Our SGM count addition method overcomes the hurdle of using sparse data matrix in CoDA. It also preserves cell-specific characteristics which can be revealed on PCA or other embedding methods. CLR shows many advantages and compatibility with various count addition methods. It also provided more distinct and biologically coherent clustering patterns in dimension reduction visualizations.

One key benefit of CoDA is its superior handling of low-quality, zero-inflated cells. By correctly identifying degraded cells as separate clusters rather than transitional states, CoDA helps prevent misinterpretation of technical artifacts as biological phenomena. We recently showed that the degraded cells were a primary cause of artefactual results and conclusions from trajectory analysis of scRNA-seq^6^. Applying CoDA to re-analyze the data will help to avoid such mistakes and wrong conclusions.

CoDA-hd is practical for routine application. Although CoDA requires more computational resources than log-normalization, the benefits in downstream analyses justify this trade-off. Furthermore, our implementation of fast truncated SVD for dimension reduction keeps CoDA practical even for large datasets. As many public dataset has been prior normalized, usually by LogNorm method, we demonstrated that LogNorm can be readily transformed by CLR to CoDA. Therefore, many public datasets can take advantage of CoDA.

It is interesting to note that CoDA LR transformations like CLR can be helpful for correct trajectory analysis. This improvement suggested that the gene ratios can be great indicators to reflect the cell continuous developing state and to describe cell-cell relationship. In biomarker prediction, biomarkers with low predictive abilities using log-normalization had much better performance when using CoDA LR transformations. The gene ratios again demonstrated their potentials to be used for classifications and clinical diagnosis.

We here demonstrate that CoDA has outstanding performance with specific count addition scheme for sparse matrices like scRNA-seq. Similar approach could also be applied to other high dimensional sparse matrices of big data in other disciplines, such as those commonly encountered in the business.

## Conclusion

The compositional nature of scRNA-seq data has been long neglected due to various reasons, like its sparse nature and lack of CoDA software. But the fact that scRNA-seq data is compositional cannot be denied. Here, we proposed theoretically sound and practical methods and associated framework to analyze high dimensional data by CoDA. By treating scRNA-seq data by CoDA, we gain certain advantages in cell clustering and visualization, handling of low-quality cells, trajectory inference, and biomarker identification.

## Abbreviation

CoDA: compositional data analysis
scRNA-seq: single cell RNA-seq
LR: log ratio
CLR: centered-log-ratio
ILR: isometric-log-ratio
TA: transcript abundances
PCA: principal component analysis
UMAP: uniform manifold approximation and projection
HTS: high-throughput sequencing
DEG: differential expressed gene
ALR: additive-log-ratio
HKGLR: housekeeping-gene-log-ratio
LRA: log-ratio analysis
SVD: singular value decomposition
ARI: adjusted rand index
NMI: normalized mutual information
PBMC: peripheral blood mononuclear cells
SCC: Spearman correlation coefficients
POS: pseudo-temporal ordering score
AUROC: area under the receiver operating characteristic curve
FC: fold changes.

## Data Availability

Public datasets used in this study were summarized in **Table S3**. Some example datasets were also placed in: https://github.com/GO3295/CoDAhd.

## Code Availability

The code for the evaluations and implementing CoDA LR transformations (**R** package ‘CoDAhd’) was placed at https://github.com/GO3295/CoDAhd.

## Notes

### Competing Interest Statement

The authors have declared no competing interest.

