## Supplemental information for "Compositional data modeling of high-dimensional single cell RNA-seq (CoDA-hd): its advantages over commonly used normalization approaches"

**Table S1. Workflow of various scRNA-seq analysis under conventional Euclidean space and CoDA**

| **Step** | **conventional analysis in Euclidean space (e.g., Seurat)** | **CoDA** |
| --- | --- | --- |
| **Implementation** | R ‘Seurat’. | e.g., R ‘easyCODA’ for conventional CoDA;  New ‘CoDAhd’ R package developed in this study. |
| **Gene counts processing** | Log-Normal transformation:  1.Feature counts for each cell are divided by the total counts for that cell.  2. Multiplied by the scale factor 10000.  3. Log transformed (add pseudocount 1).  Implemented in Seurat as NormalizeData() function. | 1. Count addition scheme for each cell (to avoid zero) (e.g., SGM).  2. Closure to 1 (counts for each cell are divided by the total counts for that cell).  3. Log-ratio transformation (e.g., CLR: each value was divided by the geometric mean of that cell, then log2 transformed). |
| **Dimension Reduction** | 1. Top 3000 variable genes were selected.  2. The log-normalized data was scaled.  3. Fast partial SVD (PCA) of the scaled log-normalized counts, implemented by Seurat.  4. UMAP of the top 10 PCs, implemented by Seurat. | 1. Top 3000 variable genes were selected.  2. The log-ratio data was scaled.  3. Partial SVD of the scaled log-ratio data, implemented by irlba package for calculation.  4. UMAP of the top 10/15 PCs, implemented by Seurat.  Alternatively, this step was implemented in easyCODA package using LRA() function (i.e., LRA refers to CLR + PCA). |
| **Clustering** | 1. Top 10 PCs were used.  2. Louvain & K-means clustering implemented by ‘scran’ and ‘igraph’ in R.  3. Evaluated by Entropy of accuracy (H_acc_), Entropy of purity (H_pur_), Adjusted Rand Index (ARI) and Normalized mutual information (NMI). | Same as conventional analysis but on log-ratio data. |
| **Trajectory inference** | 1.Performed on normalized data with default settings.  (1). Slingshot  (2). Monocle2 & 3  (3). DPT  2. Evaluated by Spearman Correlation Coefficient (SCC), pseudo-temporal ordering score (POS), false positive rate, and branch prediction performance. | Performed on log-ratio data with default settings.  Same as conventional analysis. |
| **Biomarker prediction** | 1. Select top 10 marker genes for each comparison from matched bulk comparison or ground truth of simulated datasets.  2.Evaluate their predictive performance on single cell normalized data by calculating Area Under Curve (AUC). | Same as conventional analysis but on log-ratio data. |

**Table S2. Methods evaluated in this study.**

| **Usage /Group** | **Methods** | **Platform** | **Description** |
| --- | --- | --- | --- |
| A | Log-normalization (Raw-LogNorm) (traditional method for scRNA-seq) | R | Feature counts for each cell are divided by the total counts for that cell and multiplied by the scale factor 10000. This is then log transformed (add pseudocount 1). Implemented in Seurat as NormalizeData() function. |
| A | SC-Transform (Raw-SCT) | R | Apply Seurat SCTransform() normalization (regularized negative binomial regression) and use the ‘corrected data’ assay (log space) for analyses. |
| B | MAGIC (imputation) | Python | A smoothing-based dropout (zero) imputation algorithm. We first log-normalized the data and then performed imputation. |
| B | ALRA (imputation) | R | A low-rank-matrix-based dropout (zero) imputation algorithm. We first log-normalized the data and then performed imputation. |
| CoDA | CLR | R | Centered log ratio transformation. Count addition scheme (e.g., SGM) or imputation was applied to handle the sparse matrix. Next, the data was closed to 1 for each cell (divided by the total counts for that cell, as proportion), followed by CLR (each value was divided by the geometric mean of all values (genes) in each cell. Data was then log2 transformed). |
| CoDA | ILR | R | Isometric log ratio transformation. Maps the parts of the composition in the simplex into ILR-coordinates in the Euclidean space. See Methods. |
| CoDA | HKGLR | R | Housekeeping genes log ratio transformation. Similar to CLR but use the geometric mean of the selected features (i.e., HK genes SDHA, ACTB, UBC, YWHAZ, GAPDH in this study) as denominator. |
| CoDA | MAGIC-CLR | Python | Applied the CLR transformation to the MAGIC imputed data (log space). |
| CoDA | ALRA-CLR | R | Applied the CLR transformation to the ALRA imputed data (log space). |

Nine normalization / CoDA transformation / imputation methods were used for comparison in this study. Group A are two commonly used normalization methods in Euclidean space for scRNA-seq data. Group B are two dropout imputation methods used to impute counts for dropouts in the scRNA-seq data matrices. Group CoDA are various log-ratio transformations of compositional data that have been explored on simulated and real scRNA-seq datasets to evaluate their performance.

**Table S3. Datasets used in this study.**

| **Datasets** | **Type** | **Size (Gene × Cell/Sample)** | **No. Gene with max count** $\boldsymbol{\geq}$ **10** | **Zero rate** | **Description** | **Evaluation** |
| --- | --- | --- | --- | --- | --- | --- |
| **CellBench 10x 5cl (GSE126906)** | Single cell RNA-seq | 10,164 × 3,918 | 3836 | 58.1% | Cell mixture sample with 5 cell lines H2228, H1975, A549, H838 and HCC827. | Clustering; Handle degraded cells; Biomarker prediction. |
| **GSE86337** | Bulk RNA-seq | 18,641 × 5 | - | 13.3% | Bulk samples of 5 cell lines H2228, H1975, HCC827, H838 and A549. | Biomarker prediction. |
| **GSE81861** | Single cell RNA-seq | 27,315 × 362 | 25,292 | 52.6% | Cell mixture sample with 5 cell lines A549, GM12878, H1-hESC, IMR90, and K562. | Clustering; Biomarker prediction. |
| **ENCODE bulk RNA-seq** | Bulk RNA-seq | 14,920 × 14 | - | 7.85% | Bulk samples of 5 cell lines A549, GM12878, H1-hESC, IMR90, and K562. | Biomarker prediction. |
| **GSE75748** | Single cell RNA-seq & Bulk RNA-seq | 16,606 × 997  15,607 × 755  12,352 × 19 | 16,605  15,344  - | 42.1%  44.6%  0.02% | Cell mixture sample and bulk sample with 7 cell types DEC, EC, H1, H9, HFF, NPC and TB; Time course profiling single cell sample and bulk sample using H1. | Clustering; Handle degraded cells; Pseudotime trajectory analysis; Biomarker prediction. |
| **Sorted PBMC (10x Genomics)** | Single cell RNA-seq | 8,209 × 59,620 | 1,144 | 91.2% | Sorted (known) cells from 10 cell types: B cells, CD14 monocyte cells, CD34 cells, CD4 T helper cells, CD56 NK cells, cytotoxic T cells, memory T cells, naïve cytotoxic cells, naïve T cells, and regulatory T cells) | Clustering. |
| **CellBench cellmix1-4 (GSE118704)** | Single cell RNA-seq | 6,397 × 266  9,112 × 268  9,246 × 285  11,294 × 288 | 1,093  3,054  2,986  4,149 | 70.3%  65.5%  65.0%  57.9% | 9 cell mixtures with differentiation information from three cell lines H2228, H1975 and HCC827. | Pseudotime trajectory analysis |
| **GSE79578** | Single cell RNA-seq | 14,225 × 3,331 | 2,549 | 90.2% | Mouse embryonic stem cells after different periods of continuous exposure to retinoic acid. | Pseudotime trajectory analysis |
| **GSE90047** | Single cell RNA-seq | 18,683 × 529 | 15,825 | 52.2% | Sorted hepatoblasts, hepatocytes and cholangiocytes from E10.5-E17.5 mouse fetal livers. | Pseudotime trajectory analysis |
| **Gold TI datasets (n=15)** | Single cell RNA-seq | See details in original paper  10.5281/zenodo.1443566 | | | A collection of ‘gold standard’ datasets with true time label/cell types for trajectory inference (linear type) | Pseudotime trajectory analysis |
| **Simulated dataset1** | Single cell RNA-seq (Simulated) | 10,000 × 1,500 | 5,450 | 93.0% | Splatter: splatEstimate using CellBench 10x 5cl; nGenes=10000, batchCells=1500, group.prob=c(0.3,0.25,0.2,0.15,0.1); set dropout.mid to let dropout rate=92%. Others are default. | Clustering; Biomarker prediction. |
| **Simulated dataset2** | Single cell RNA-seq (Simulated) | 10,000 × 1,500 | 6,699 | 75.7% | Splatter: splatEstimate using CellBench 10x 5cl; nGenes=10000, batchCells=1500, group.prob=c(0.3,0.25,0.2,0.15,0.1); set dropout.mid to let dropout rate=68%. Others are default. | Clustering; Biomarker prediction. |
| **Simulated dataset3** | Single cell RNA-seq (Simulated) | 10,000 × 1,500 | 6,825 | 66.0% | Splatter: splatEstimate using CellBench 10x 5cl; nGenes=10000, batchCells=1500, group.prob=c(0.3,0.25,0.2,0.15,0.1); set dropout.mid to let dropout rate=43%. Others are default. | Clustering; Biomarker prediction. |
| **Simulated dataset4** | Single cell RNA-seq (Simulated) | 8,000 × 1,500 | 5,395 | 90.1% | SplatPop: nGenes=8000, batchCells=200, group.prob=c(0.4,0.3,0.3), condition.prob=c(0.5,0.5), similarity.scale=15, de.facLoc=0.5, de.facScale=0.5, cde.facLoc=0.38, cde.facScale=0.38, set dropout.mid to let dropout rate=90%. Others are default. | Clustering. |

Twenty-nine real datasets (including a collection of 15 ‘gold standard’ trajectory datasets) and four simulated datasets (i.e., Splatter and SplatPop) were used for different downstream analyses.

**Figure S1. Illustration of Compositional data analysis (CoDA) for different high-throughput sequencing data**

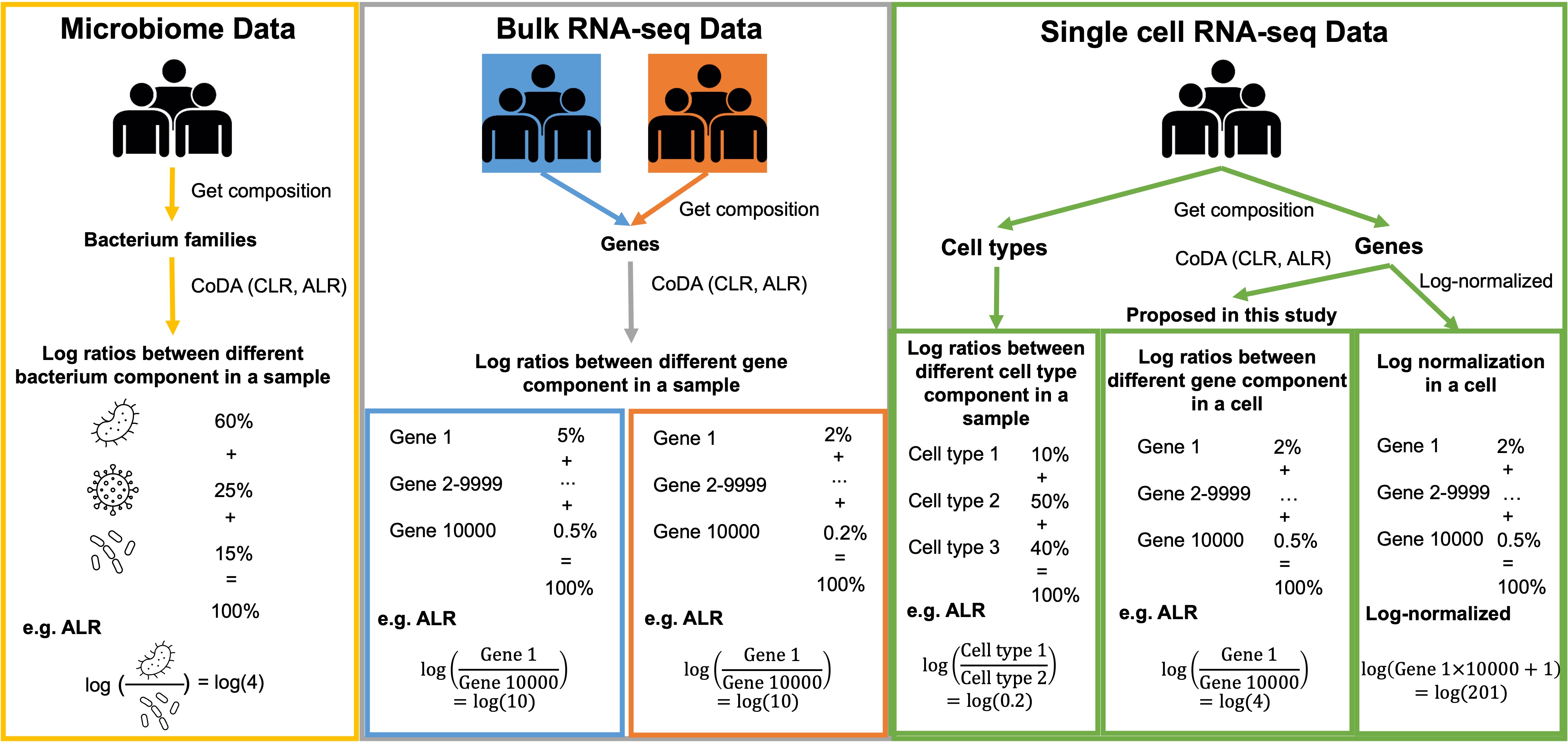

**Figure S2. 2-D PCA & UMAP plots and elbow plots of CLR+partial SVD and LRA (easyCODA) using the CellBench-10X-5CL dataset (top 3,000 features×3,918 cells).**

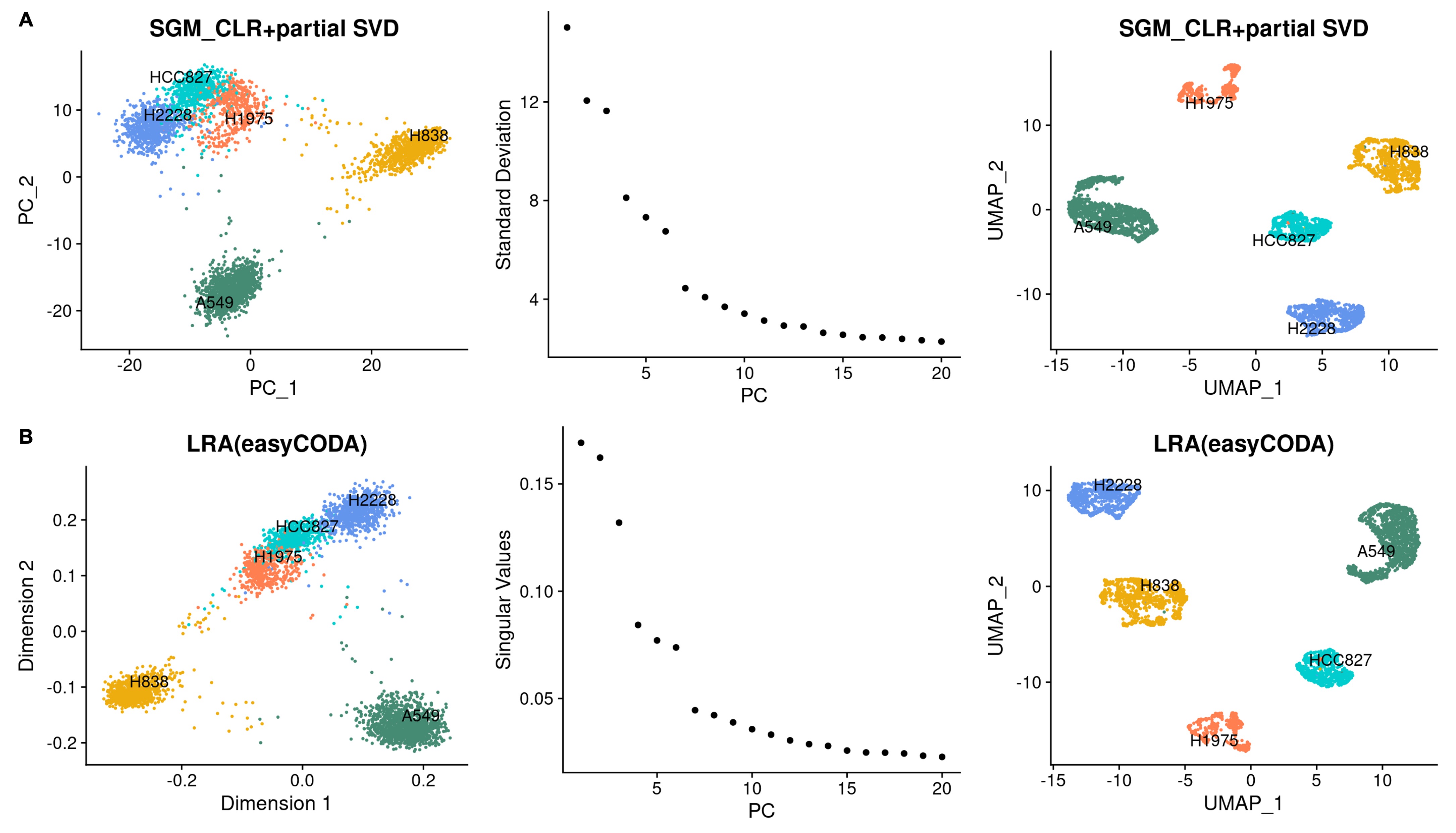

**Figure S3. Clustering performance of different count additions in CoDA transformations using K-means and Louvain Clustering algorithms on real and simulated datasets (Raw-LogNorm as baseline) (individual dataset).**

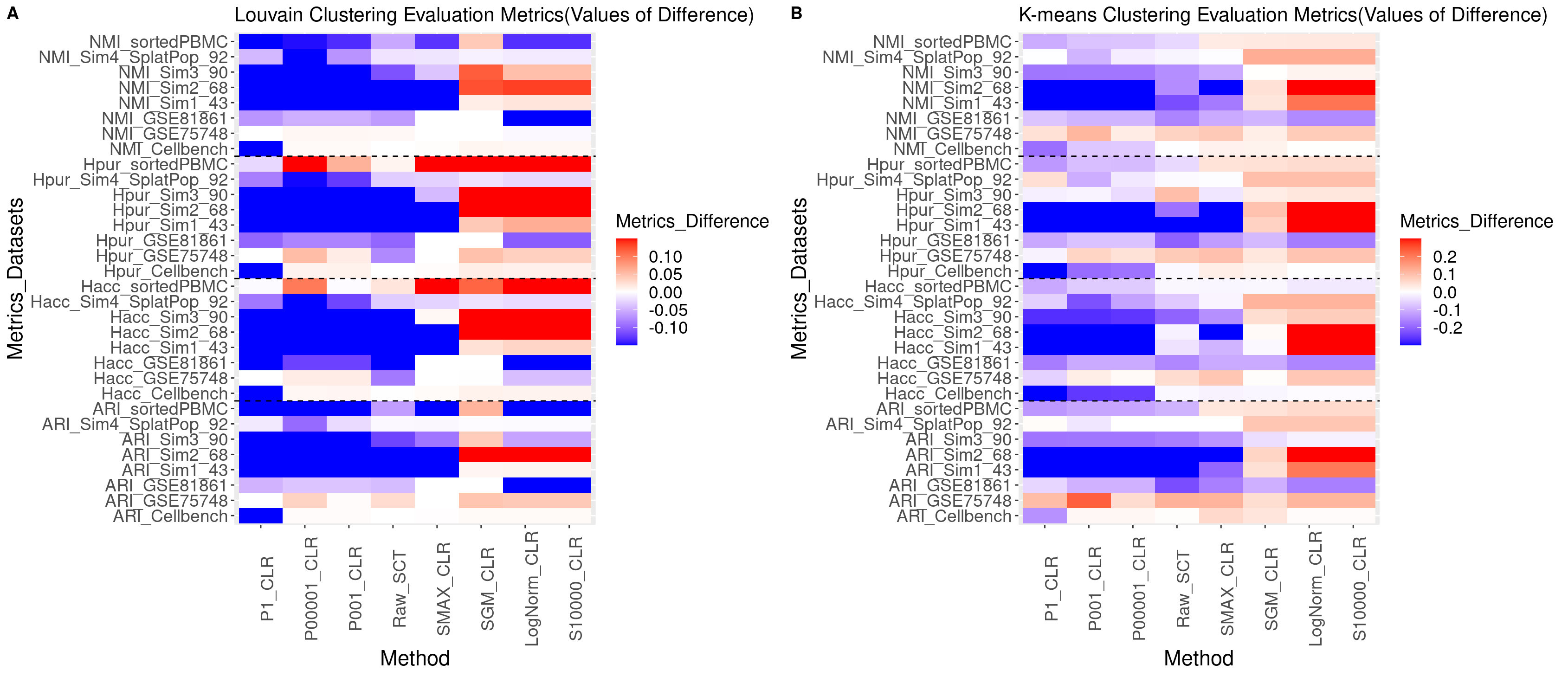

**Figure S4. Clustering performance of different normalizations and CoDA transformations using K-means and Louvain Clustering algorithms on real and simulated datasets (Raw-LogNorm as baseline) (individual dataset)**

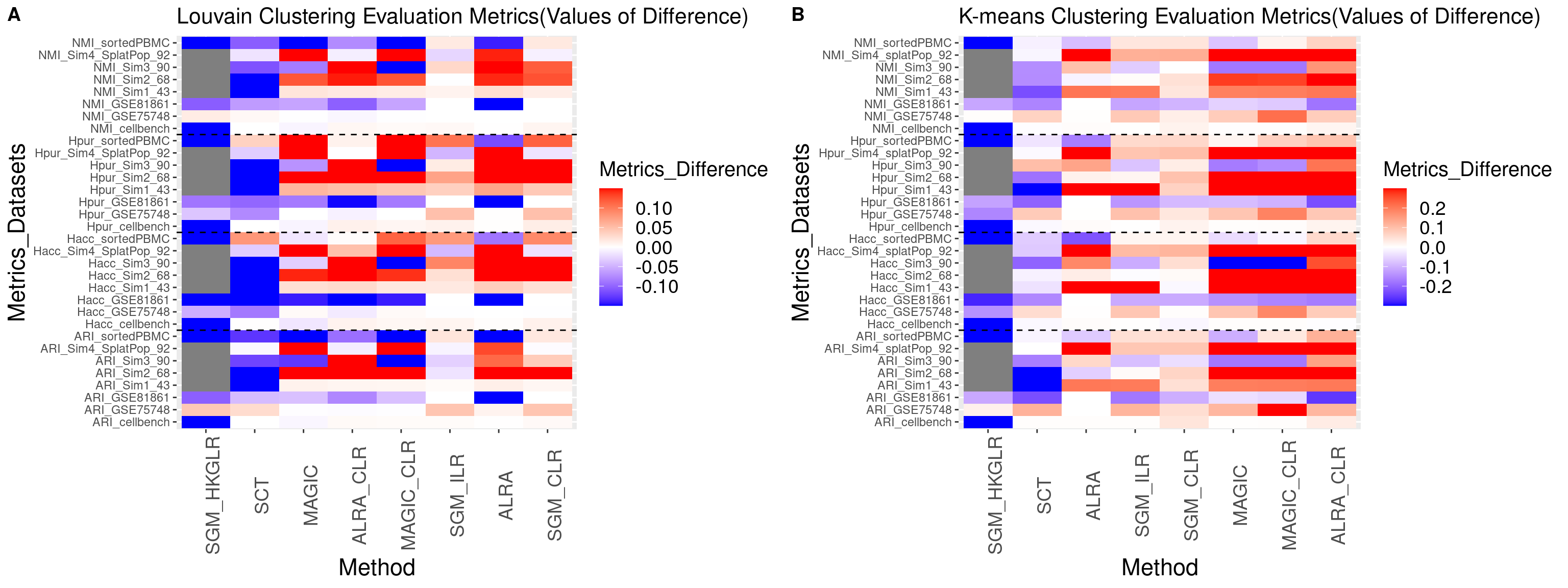

**Figure S5. Clustering performance of different count additions in CoDA transformations using K-means and Louvain Clustering algorithms on simulated ground truth datasets (True-LogNorm as baseline)**

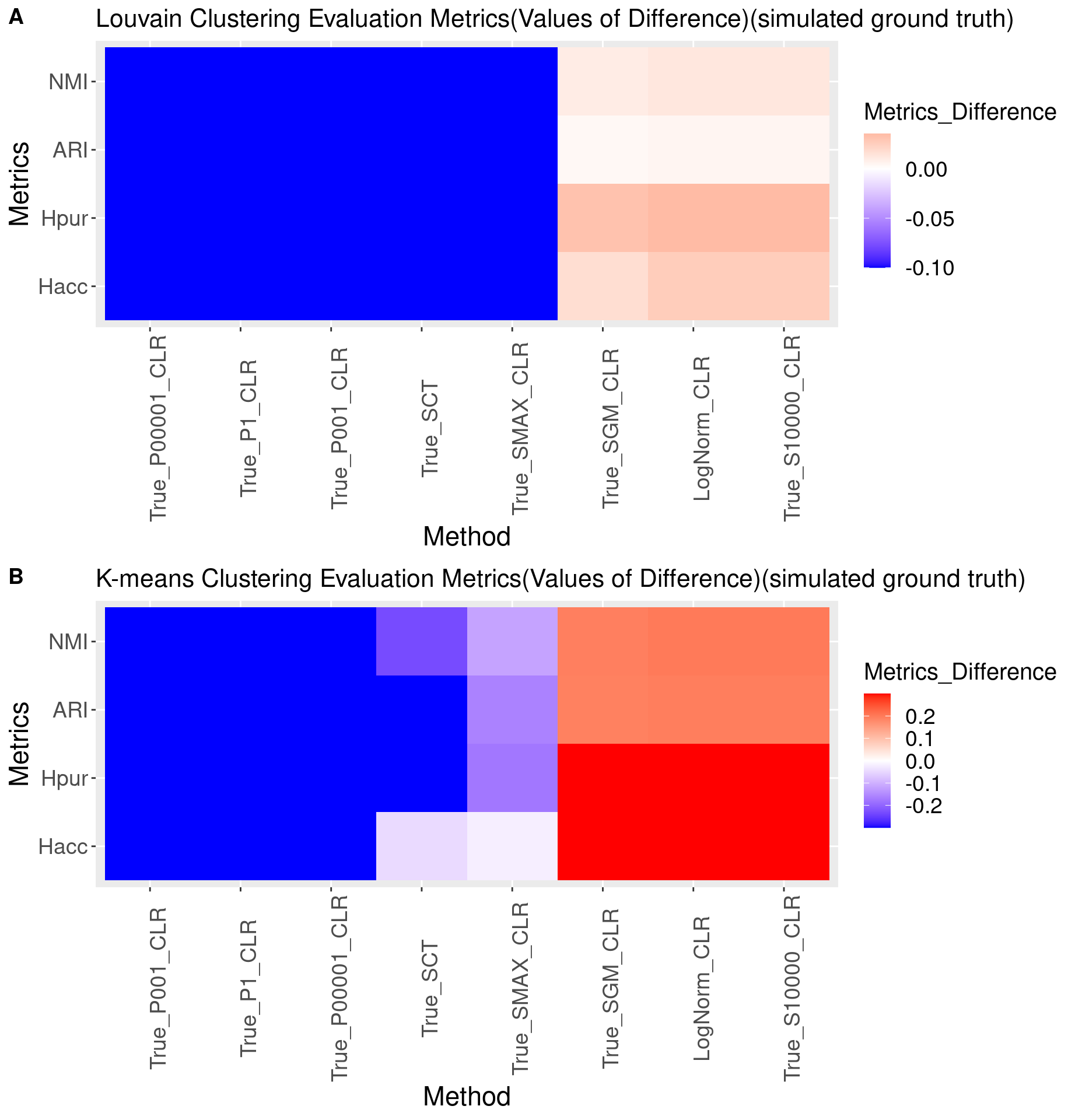

**Figure S6. 2-D UMAP plots of different CoDA count additions using the sorted PBMC dataset**

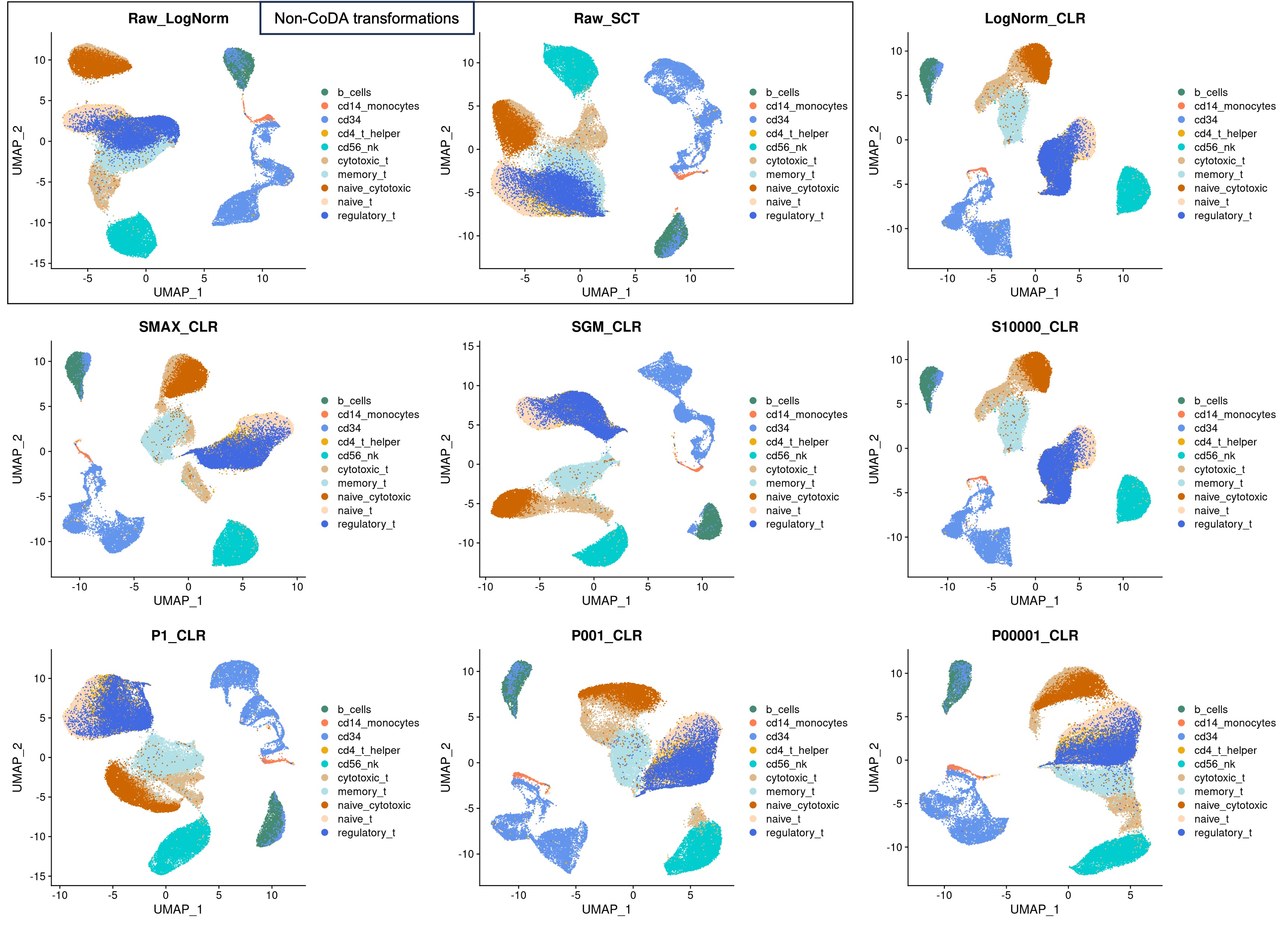

**Figure S7. 2-D UMAP plots of different normalizations and CoDA transformations using the sorted PBMC dataset**

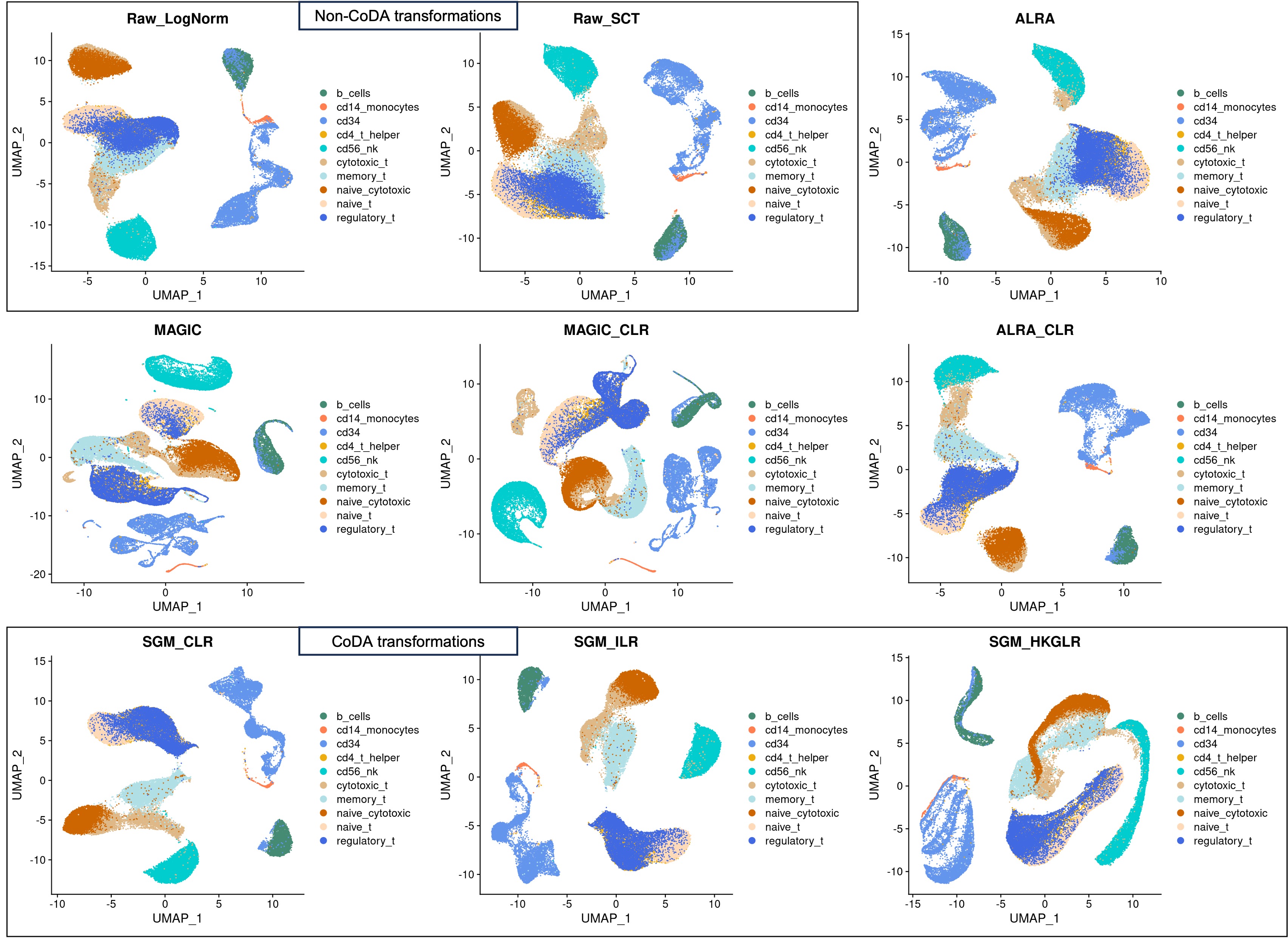

**Figure S8. 2-D PCA plots of different CoDA count additions using the CellBench-10X-5CL dataset**

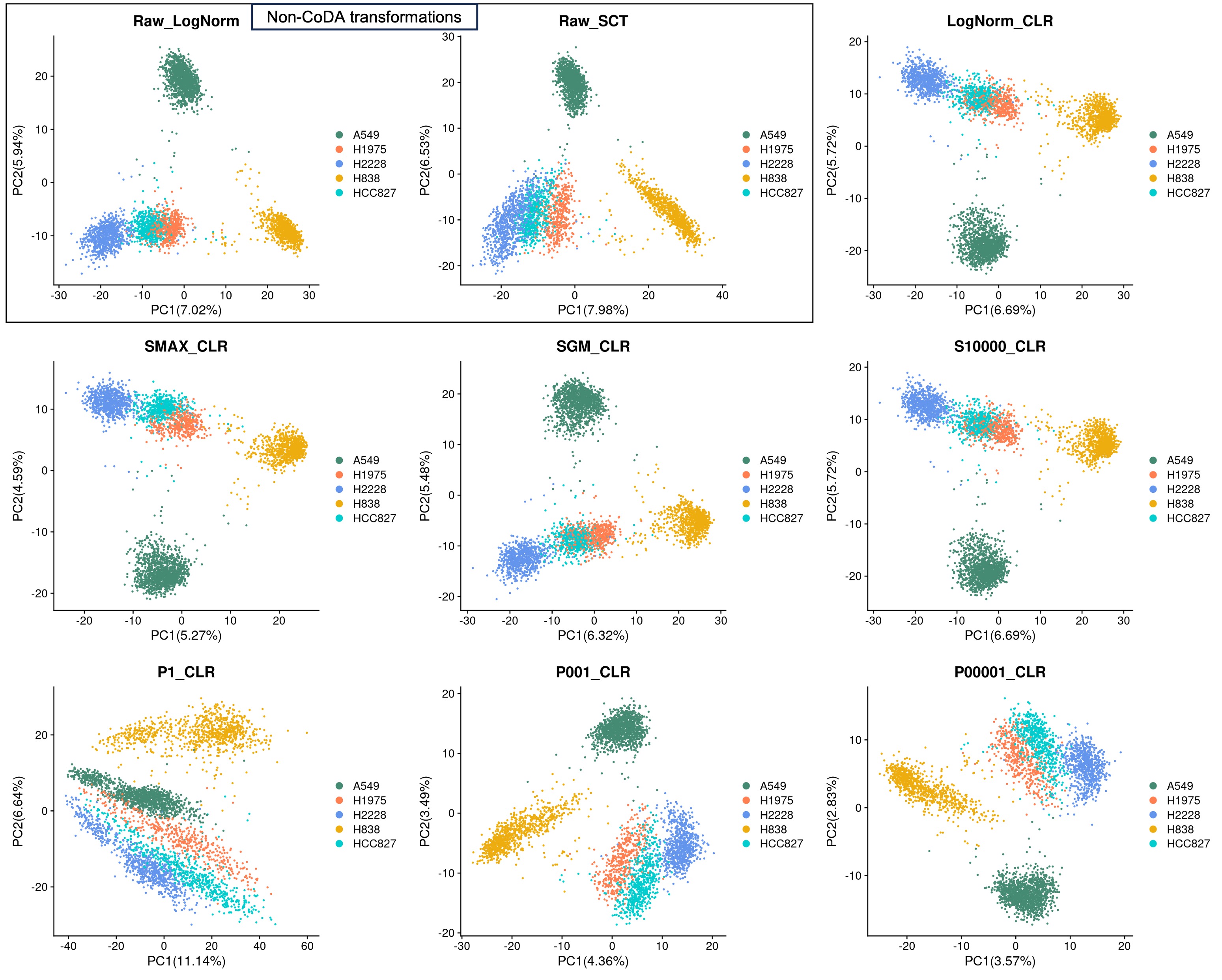

**Figure S9. 2-D UMAP plots of different CoDA count additions using the CellBench-10X-5CL dataset**

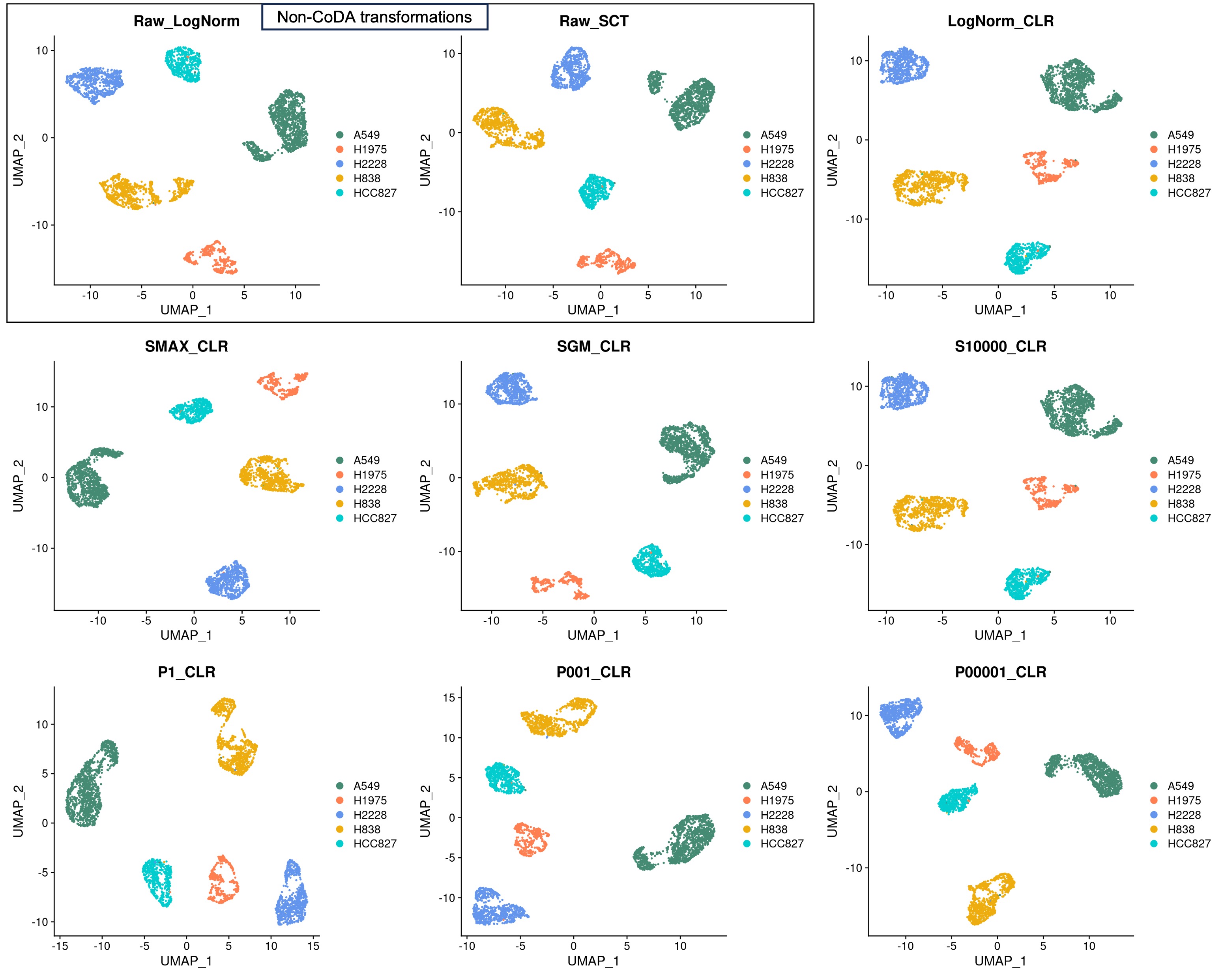

**Figure S10. 2-D PCA plots of different normalizations and CoDA transformations using the CellBench-10X-5CL dataset**

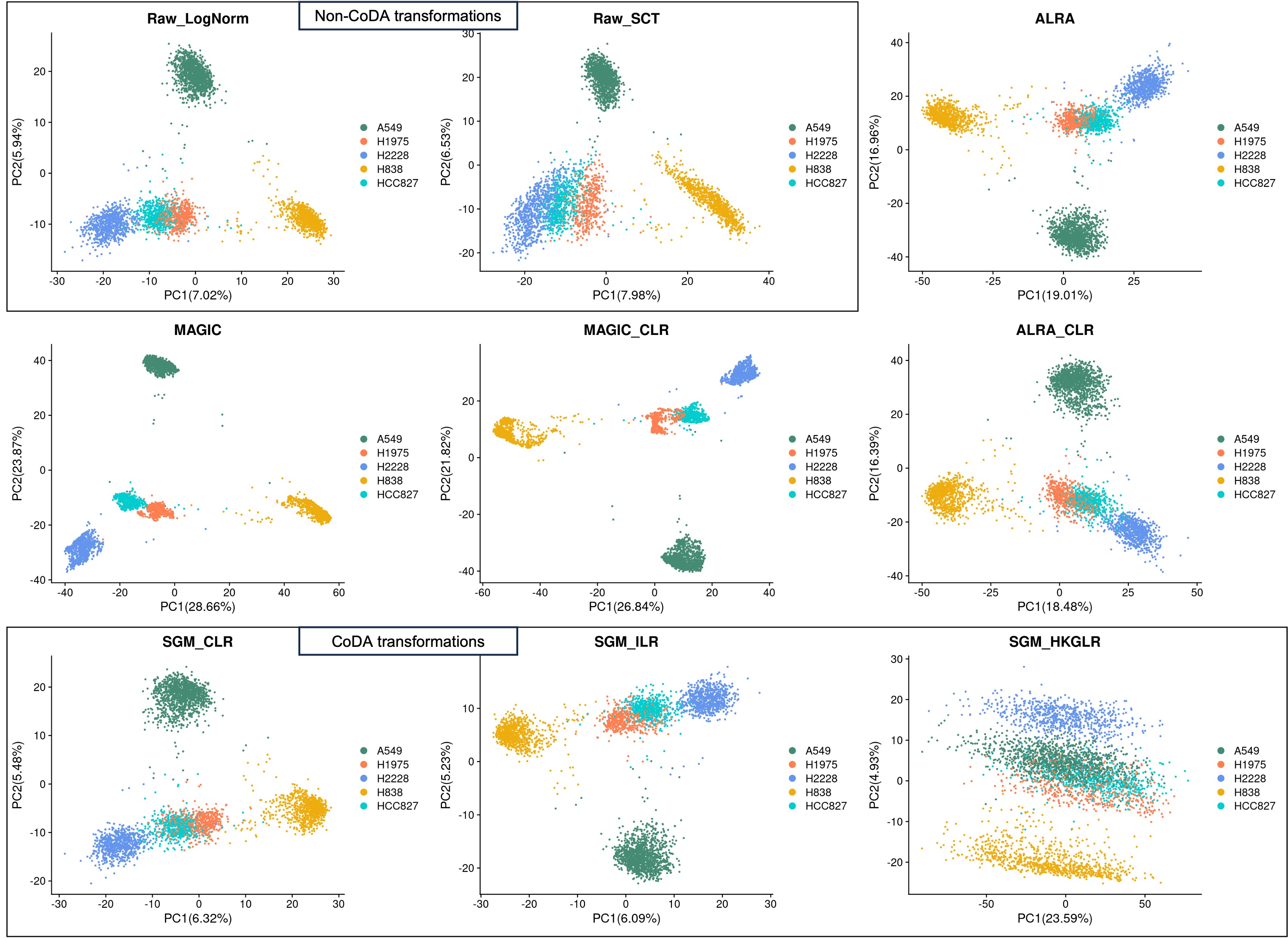

**Figure S11. 2-D UMAP plots of different normalizations and CoDA transformations using the CellBench-10X-5CL dataset**

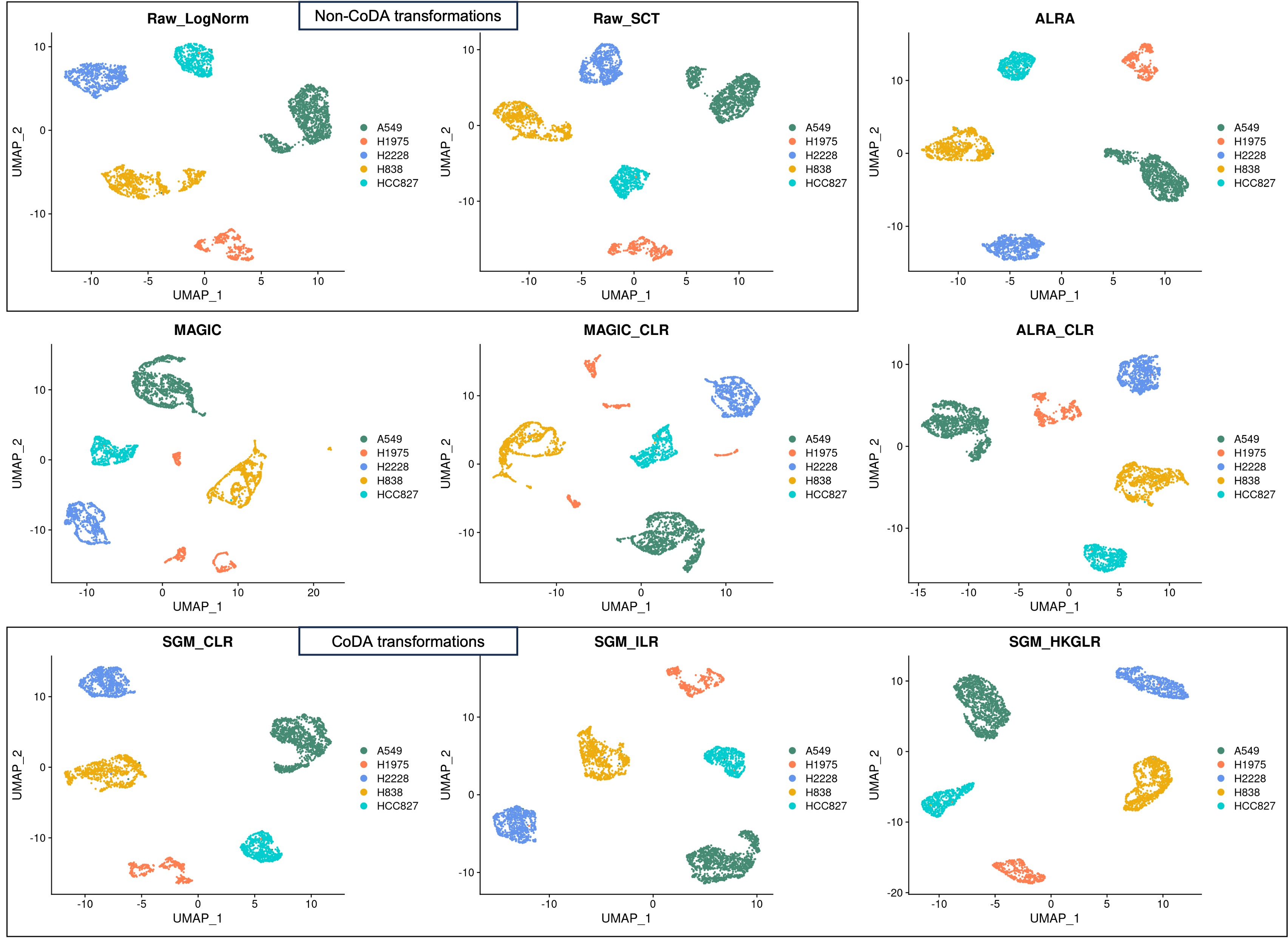

**Figure S12. 2-D PCA plots of different normalizations and CoDA transformations using the simulated dataset 2**

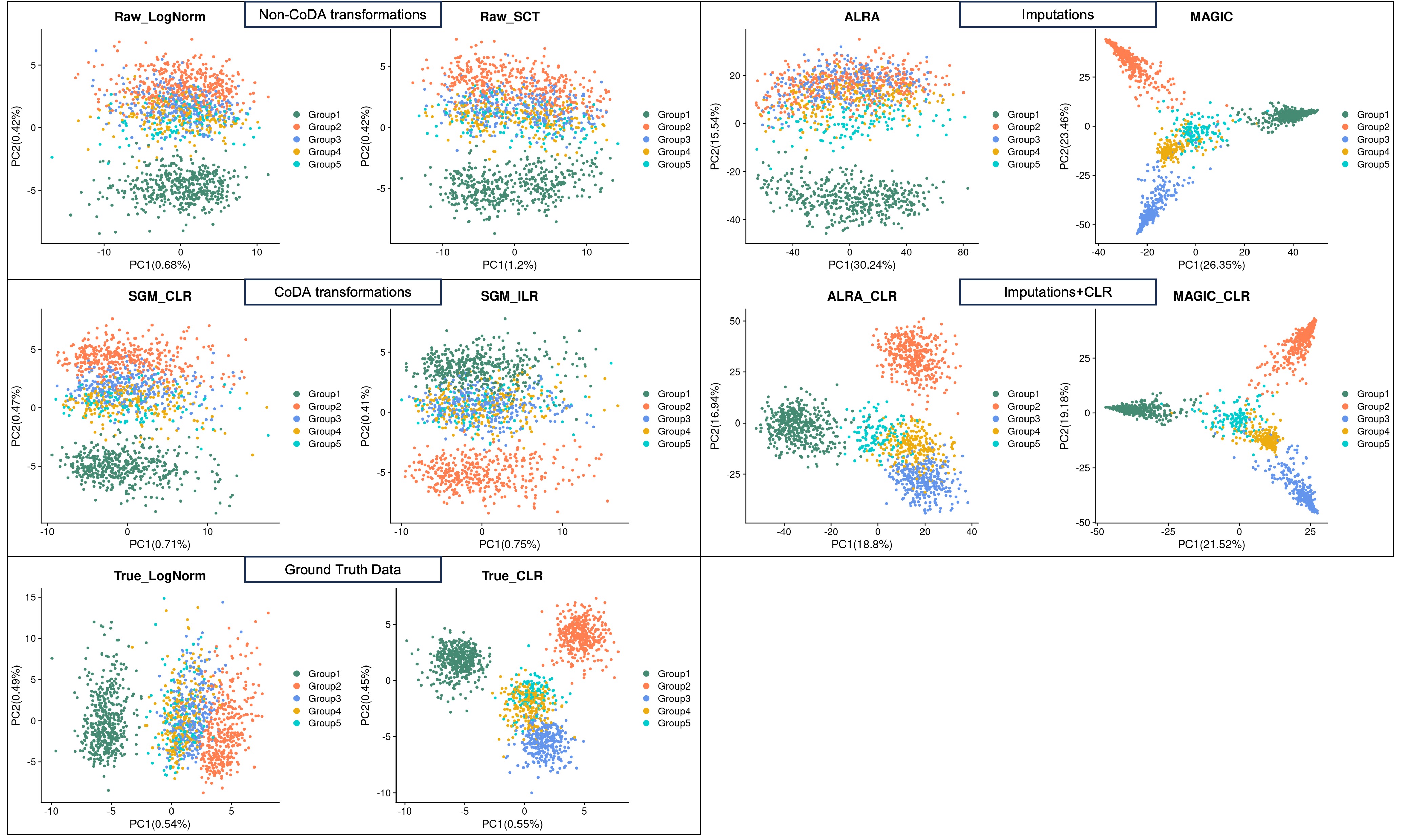

**Figure S13. 2-D UMAP plots of different normalizations and CoDA transformations using the simulated dataset 2**

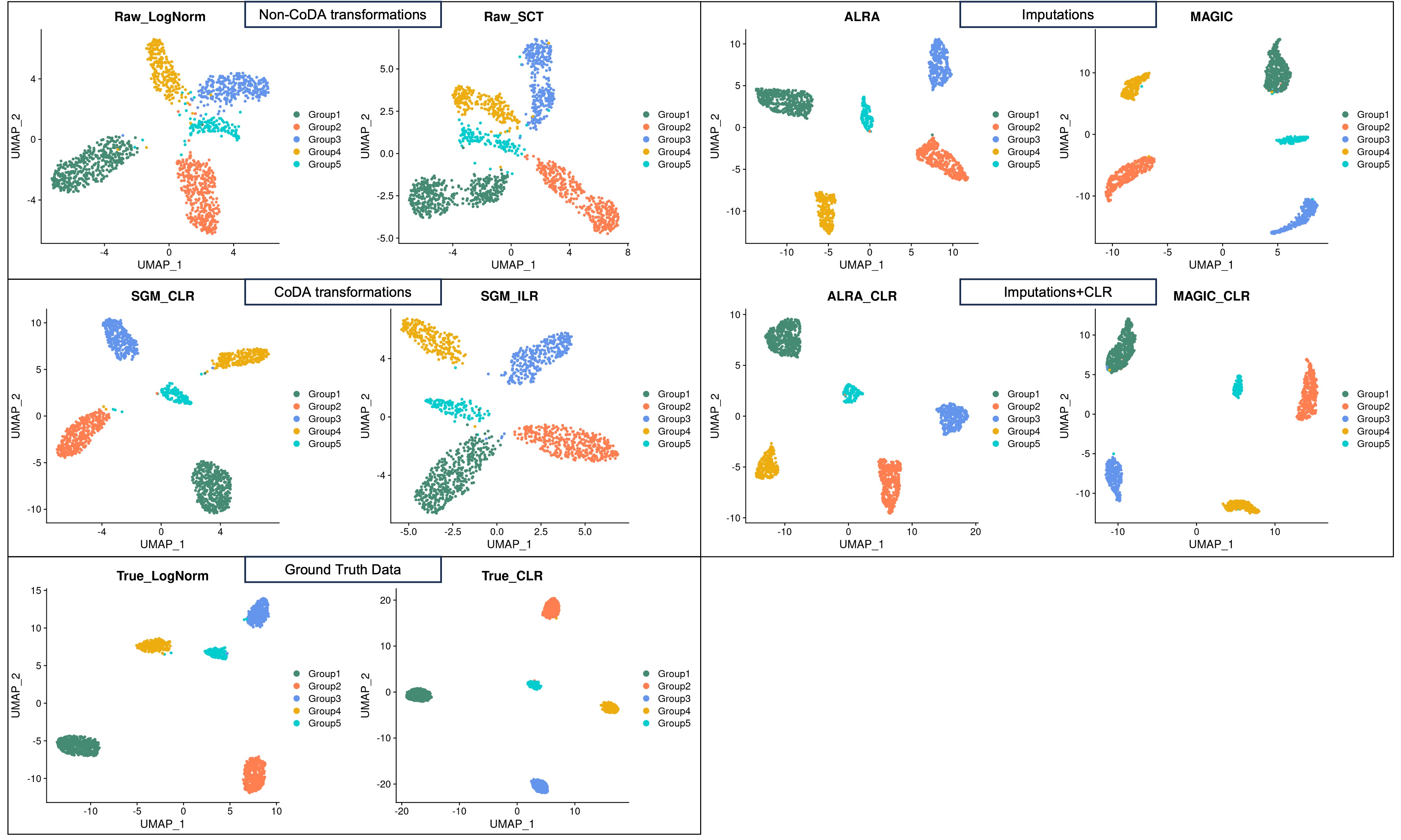

**Figure S14. 2-D PCA plots of different methods using the subset CellBench-10X-5CL dataset with random-zeros-H1975-cells and the subset GSE75748-CellType dataset with random-zeros-H1-cells.**

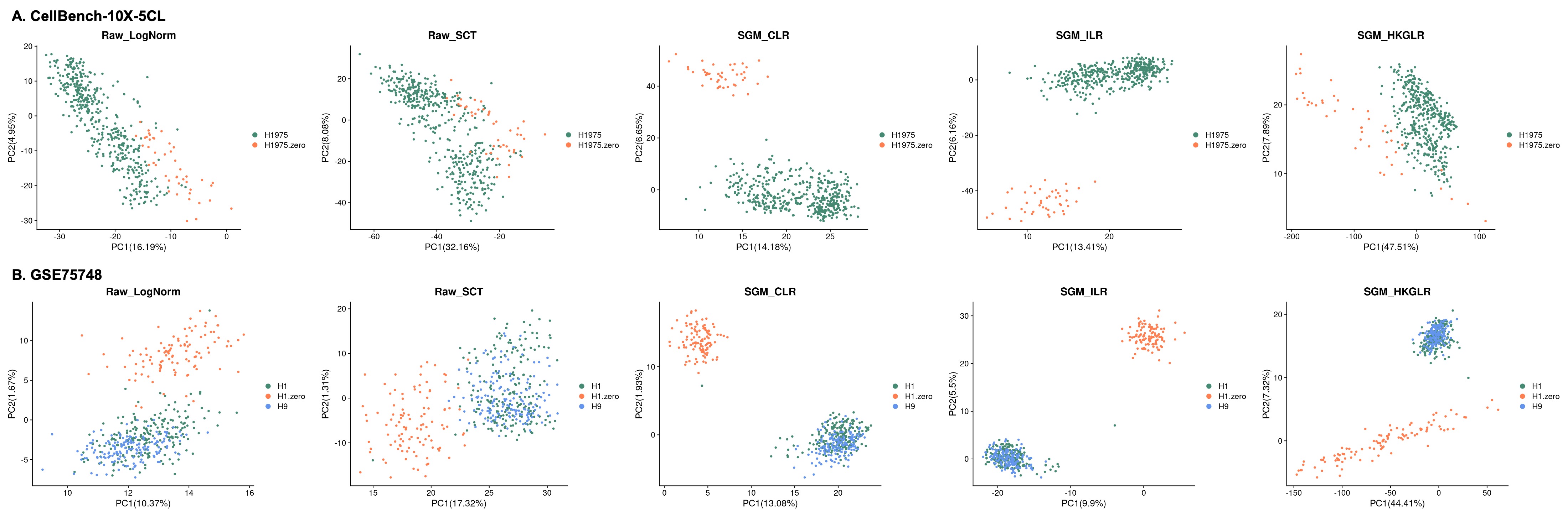

**Figure S15. Performances of different normalizations and CoDA transformations on Slingshot pseudotime trajectory analysis in each of the datasets (Raw-LogNorm as baseline).**

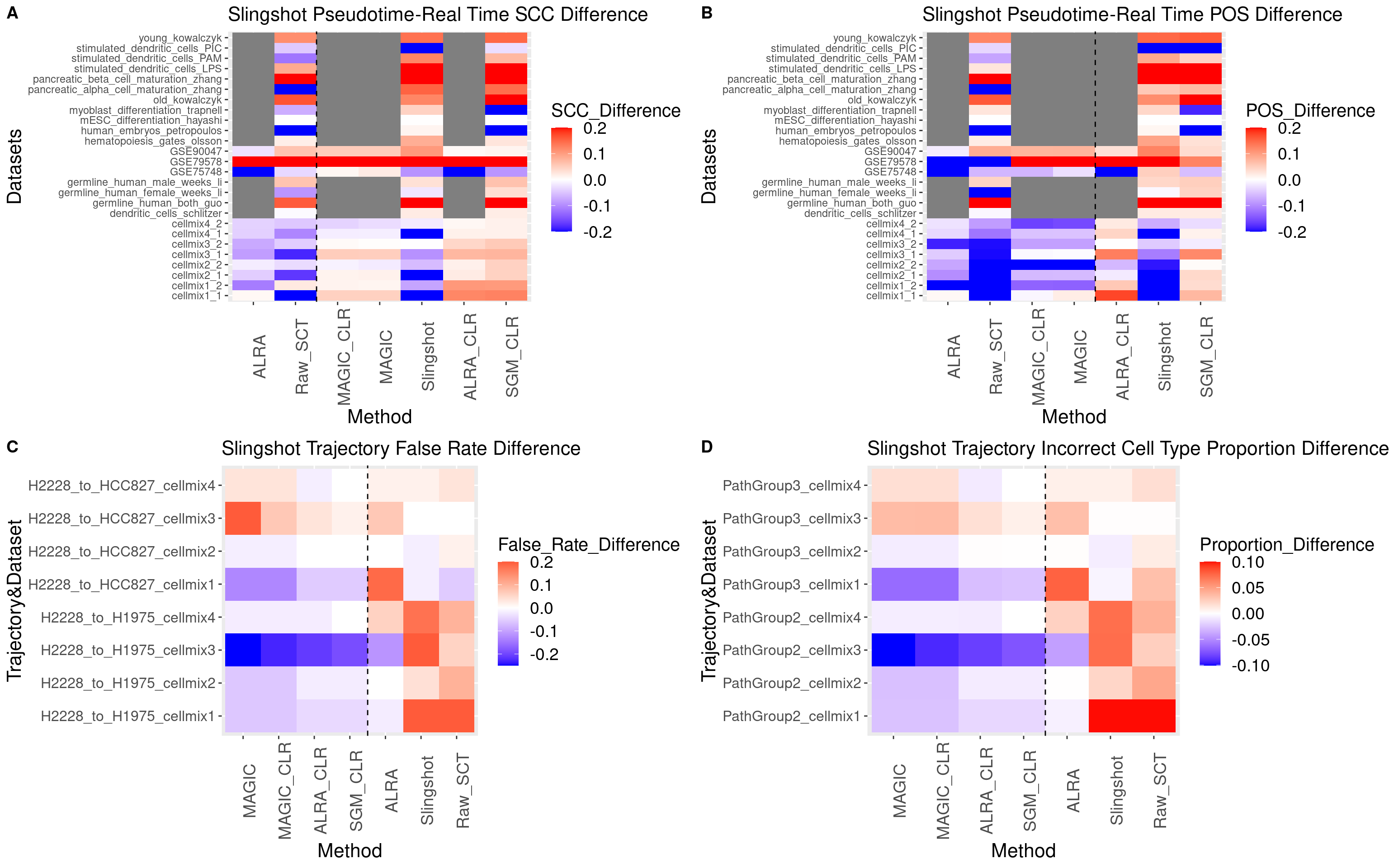

**Figure S16. Performances of different normalizations and CoDA transformations on DPT pseudotime trajectory analysis.**

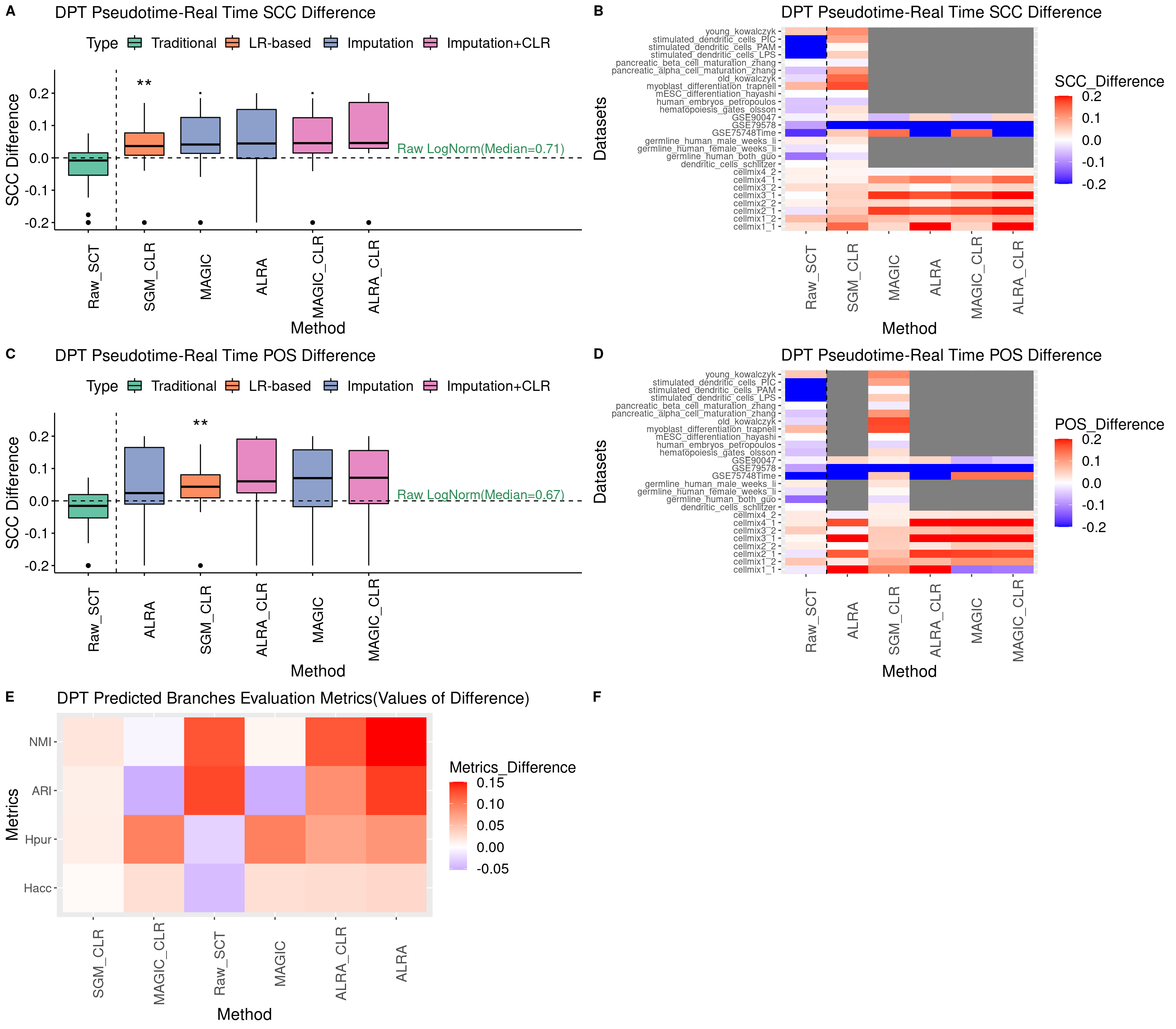

**Figure S17. Performances of different normalizations and CoDA transformations on Monocle2 & 3 pseudotime trajectory analysis.**

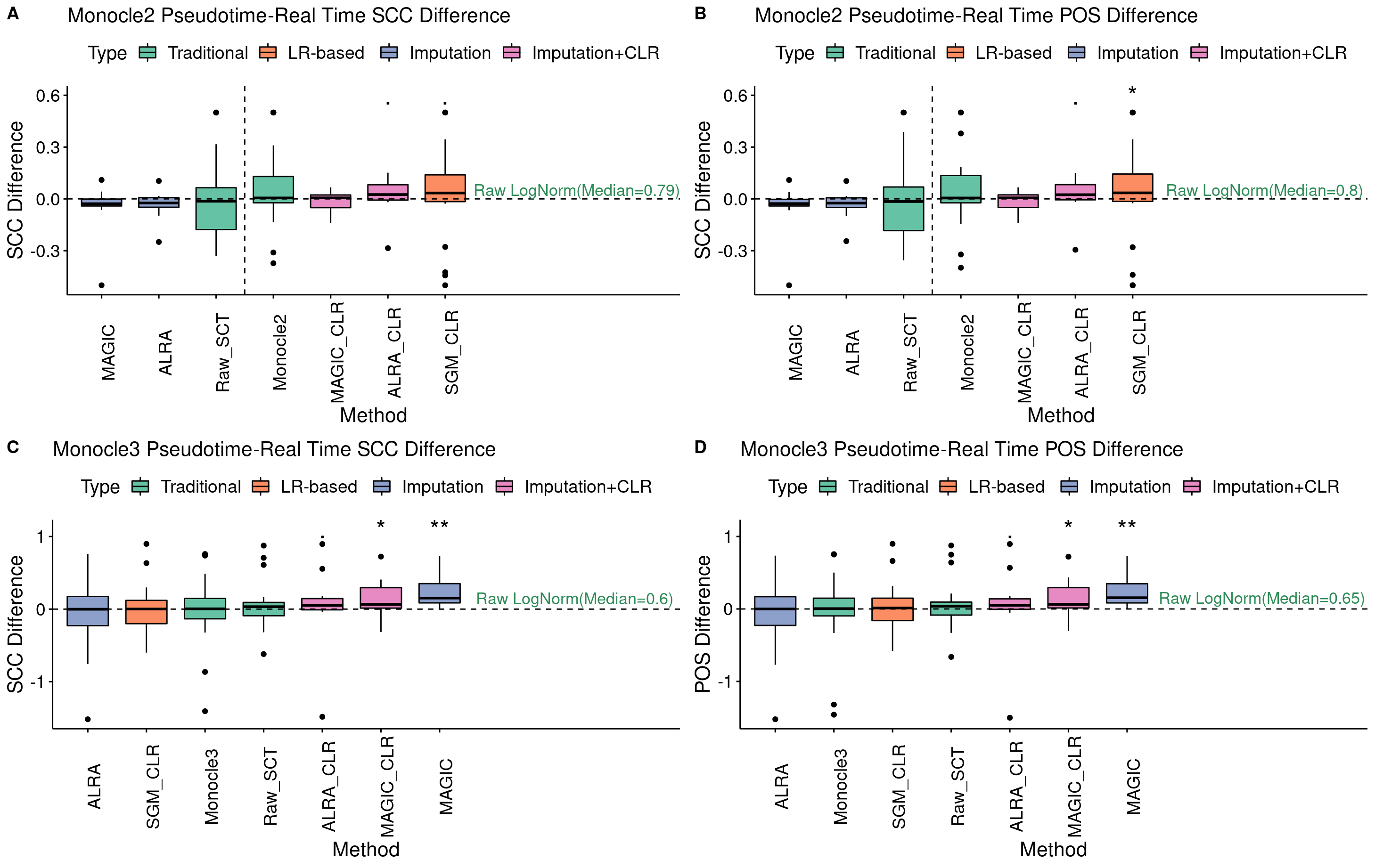
